# Computational Modeling of the Anti-Inflammatory Complexes of IL37

**DOI:** 10.1101/2024.09.21.613817

**Authors:** Inci Sardag, Zeynep Sevval Duvenci, Serkan Belkaya, Emel Timucin

## Abstract

Interleukin (IL) 37 is an anti-inflammatory cytokine belonging to the IL1 protein family. Owing to its pivotal role in modulating immune responses, particularly through interfering with the IL18 signaling, elucidating the IL37 complex structures holds substantial therapeutic promise for various autoimmune disorders and cancers. Although the structural homology between IL37 and IL18 suggests a common binding mechanism with the primary members of IL18 signaling, the structures of IL37 complexes have not been experimentally resolvet yet. This computational study aims to address this gap through molecular modeling and classical molecular dynamics simulations, revealing the structural underpinnings of its modulatory effects on the IL18 signaling pathway. All IL37 protein-protein complexes, including both receptor-dependent and receptor-independent pairs, were modeled using a range of methods from homology modeling to AlphaFold2 multimer predictions. The models that successfully captured experimental features were subjected to molecular dynamics simulations. As positive controls, binary and ternary PDB complexes of IL18 were also included. The comparative look on the IL37 and IL18 complexes revealed a highly dynamic nature for the IL37 complexes. Repeated simulations of IL37-IL18Rα showed altered receptor conformations capable of accommodating IL37 in its dimeric form without clashes, providing a structural basis for the failure of IL18Rβ to be recruited to the IL37-IL18Rα complex. Simulations of receptor complexes involving various mature forms of IL37 revealed that the N-terminal loop of IL37 is pivotal in modulating receptor dynamics. Additionally, the glycosyl chains on the primary receptor residue N297 act as a steric block against the IL37’s N-terminal loop. The interactions between IL37 and IL18BP were also investigated, and our dynamical models indicated that a homologous binding mode was unlikely, suggesting an alternative mechanism by which IL37 functions as an anti-inflammatory cytokine upon binding to IL18BP. Altogether this study accesses to the structure and dynamics of IL37 complexes, offering molecular insights into IL37’s inhibitory function within the IL18 signaling pathway and informing future experimental research.

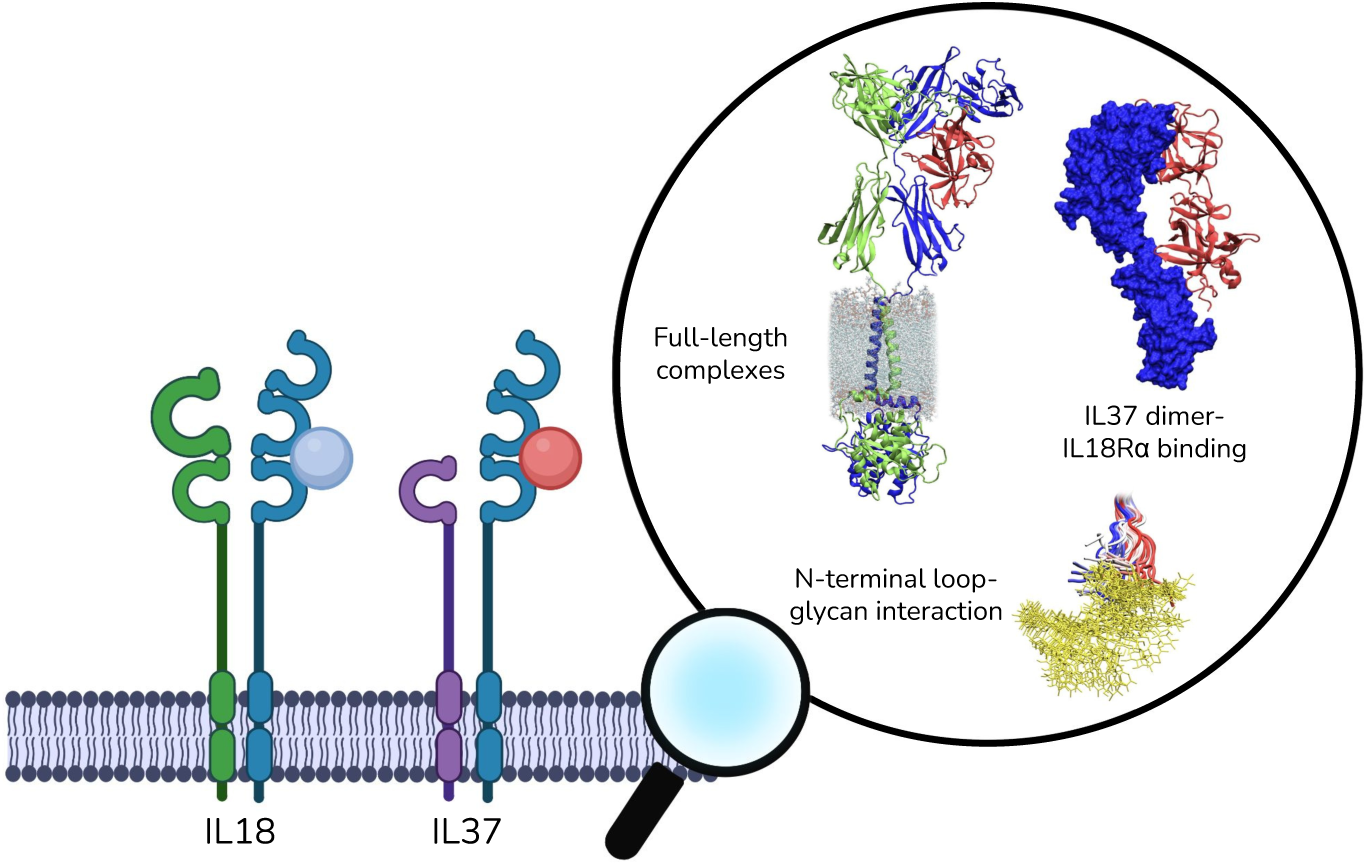
Graphical Abstract.

## 1. Introduction

The interleukin-1 (IL1) protein family plays a crucial role in mediating innate immunity and shaping the outcome of inflammation [1, 2, 3]. While most members of this family induce a proinflammatory response, a few have been shown to exert anti-inflammatory effects. Among the latter, IL37 is known to produce an anti-inflammatory effect by downregulating proinflammatory signals both extracellularly and intracellularly [4].

IL18 signaling machinery is particularly essential for the extracellular anti-inflammatory capacity of IL37. Within this machinery, IL18, another member of IL1 family, initiates a proinflammatory signal by binding to its receptor α (IL18Rα). The binary IL18-IL18Rα complex then recruits the accessory receptor (IL18Rβ), forming the ternary signaling complex [5, 6, 7]. The intracellular region of this ternary complex next engages with the adapter molecules of myeloid differentiation factor 88 (MyD88) to propagate the signal inside the cell [8]. This propagation can be disrupted at the binary complex level by IL18 binding protein (IL18BP) that sequesters IL18 from the extracellular space, preventing its binding to IL18Rα [7, 2, 9]. IL37 has been shown to hijack both of these primary partners of IL18, which are IL18Rα and IL18BP [10]. Essentially, IL37 can bind to IL18Rα, which impairs the recruitment of the IL18Rβ and blocks the formation of the ternary signaling complex [11]. Instead, the IL37-IL18Rα complex recruits an orphan receptor, IL1R8 (SIGIRR), that decoys the binding of MyD88 [8]. The complexes formed by IL37 have been reported to have weaker affinity than those formed by IL18. Specifically, recombinant IL37 exhibited ∼50 times lower affinity towards IL18Rα than IL18 [12]. IL37-IL18BP interaction was also reported to be weaker than that of IL18-IL18BP [11]. Furthermore, IL37 has also been shown to bind to Smad3 intracellularly, forming the IL37-Smad3 complex that interferes with STAT activation and inhibits inflammation [13, 14]. Altogether, IL37’s extracellular anti-inflammatory effects depend on the interplay between IL18Rα, IL1R8, and IL18BP, whilst Smad3 is a key partner for its receptor-independent intracellular role.

Earlier findings identifying IL37 complexed with components of the IL18 signaling system [4], along with the significant structural resemblance between these two cytokines [15], suggested that IL37 would adopt a similar binding mechanism to IL18 in their respective complexes. In line with this suggestion, a previous homology model first captured the IL37-IL18Rα complex, exhibiting the same binding mode with IL18 [11]. Nevertheless, no other experimental or computational structures have yet resolved the atomic details of any IL37 complexes involved in receptor-dependent or receptor-independent pathways. Given the extensive role of IL37 in both innate and adaptive immunity, elucidating the structural details of IL37 complexes holds significant therapeutic potential for a wide range of diseases, including autoimmune diseases and cancer. This end prompted this computational study to model a comprehensive set of IL37 complexes capturing experimental features and to particularly understand the structural details of how IL37 hijacks the IL18 signaling machinery. We analyzed the comprehensive set of IL37 models including the receptor-dependent and –independent complexes through molecular modeling, all-atom simulations and binding free energy predictions. Our findings unveil structural details of the IL37’s antiinflammatory action, underlining the interplay between the N-terminus of IL37 and one of the glycosylated receptor amino acids for IL18Rα interactions.

## 2. Methods

### 2.1. IL37 Structures

Homodimer structure of IL37 (PDB ID 5hn1) was retrieved based on our previous research on the stability of dimeric and monomeric IL37 [16] and its chain A, which has shorter unmodeled regions compared to the chain B, was used for modeling [15]. Internal missing regions in this structure was completed by loop modeling and the resulting structure served as the most dominant isoform, IL37b spanning the region between 49-206. Next, we also generated IL37 structures whose N-terminus was either shorter (53-206, 57-206) or longer (21-206) than the IL37b structure. Ten predictions were performed for each loop modeling and final models were selected based on lowest zDOPE scores [17]. Loop modeling was performed using MODELLER plugin [18] of ChimeraX [19].

### 2.2. Generation of IL37 Complexes

We modeled eleven IL37 complexes that were associated with an anti-inflammatory response. For complex modeling, homology modeling, proteinprotein docking and AF2 multimer predictions were conducted. For homology modeling, we adapted the IL18 binding mode in the final complexes. The resulting complexes were examined for steric clashes using UCSF ChimeraX [19] and analyzed against experimental data. Only complexes that met the criteria of experimental observations were selected. A summary of the predicted complexes were given in Table 1, including both receptor-dependent and –independent complexes. For the complexes that were generated relying on IL18 homology, we have also recruited the PDB complexes of IL18 as controls to our study. These included the binary (PDB ID: 3wo3), ternary signaling complexes (PDB ID: 3wo4).

**Table 1:** Models Generated in This Study.

|  | **Cytokine<sup>†</sup>** | **Partner (source)** | **Complex Source** |
| --- | --- | --- | --- |
| monomer | IL37 (21-206) | - | PDB (5hn1:A)/MODELLER |
|  | IL37 (49-206) | - | PDB (5hn1:A) |
| soluble<br>binary<br>receptor<br>complexes | IL18 | IL18Rα | PDB (3wo3) |
| | IL37 (49-206) | IL18Rα (3wo3) | IL18 homology |
| | IL37 (49-206) | IL18Rα (3wo4) | IL18 homology |
| | IL37 (53-206) | IL18Rα (3wo4) | IL18 homology |
| | IL37 (57-206) | IL18Rα (3wo4) | IL18 homology |
| soluble<br>ternary<br>receptor<br>complexes | IL18 | IL18Rα-Rβ | PDB (3wo4) |
| | IL37 (49-206) | IL18Rα-Rβ | IL18 homology |
| | IL37 (49-206) | IL18Rα-short Rβ | IL18 homology |
| | IL37 (49-206) | IL18Rα-IL1R8 | IL18 homology |
| full-length<br>receptor<br>complexes | IL18 | IL18Rα | IL18 homology/AF2 |
| | IL18 | IL18Rα-Rβ | IL18 homology/AF2 |
| | IL37 (49-206) | IL18Rα | IL18 homology/AF2 |
| | IL37 (49-206) | IL18Rα-IL1R8 | IL18 homology/AF2 |
| other<br>complexes | IL37 (49-206) | IL18BP | IL18 homology |
|  | IL37 (49-206) | Smad3 | AF2 |

<sup>†</sup>Chain A of the 5hn1 structure was the source of IL37 in all complexes.

#### 2.2.1. AlphaFold2 Predictions

AlphaFold2 multimer v3 (AF2) and ColabFold were used to predict IL37 complex structures [20, 21, 22]. For all predictions, MSA mode was selected as mmseqs2 uniref env while pair mode was selected as unpaired paired [23]. The number of cycles was adjusted to six and top-ranking model was relaxed by AMBER force field in 200 iterations [24]. For prediction of the full-length complexes including the toll-interleukin-1 receptor (TIR) domains, the soluble ternary complex of IL18 in PDB (3wo4) was used as the template [5]. Transmembrane AlphaFold database (TmAlphaFold) was used to determine the membrane localization of the receptors [25]. Terminal loops of all receptors with relatively low pLDDT scores (<50) were not included in the final models.

#### 2.2.2. Protein-protein Docking

ClusPro using FFT-based Piper clustering algorithm [26, 27, 28, 29] was implemented for rigid-body docking of IL37 and receptor complexes. For the IL37 structure, the chain A of the 5hn1 PDB structure was used and for the IL18 receptor α, the chain B of the PDB structure with ID 3wo3 was used. Top-scoring ten models were analyzed. Flexible docking was performed using HADDOCK 2.4, with the search space restricted to the IL18Rα residues at the interface of the binary PDB complex (3wo3) and the IL37 residues that are homologous to IL18 in this complex [30, 31].

### 2.3. System Preparation

Structures that were directly obtained from the PDB or modeled in this study were analyzed by all-atom molecular dynamics (MD) simulations. Crystal waters and other buffer contaminants were removed from the structures, while N-linked glycans were maintained in the IL18Rα, IL18Rβ and IL18BP according to their PDB information. Disulfide bonds between 22-41, 43-81, 119-158, 140-185, 237-298 in IL18Rα, 46-126, 155-180, 175-221, 273-337 in IL18Rβ, and 64-89, 86-150, 131-74 in IL18BP were also introduced according to the PDB information. The structures were then positioned at the center of orthorhombic boxes, ensuring a distance of 10 Å from the macromolecules in all directions. After solvation, all systems were neutralized to a final concentration of 150 mM using the counterions of Na^+^ and Cl^−^. All system preparation steps were performed with CHARMM-GUI [32, 33, 34].

Full-length (transmembrane) complexes were embedded in a lipid bilayer composed of phosphatidylcholine (POPC) by CHARMM-GUI bilayer builder [35, 36, 37, 38, 39]. Complexes were positioned within the bilayer membrane using the PPM 2.0, which is based on Orientations of Proteins in Membranes (OPM) database [40]. All other parameters and procedures for system preparation including protonation, solvation and neutralization were carried out as described for soluble complexes.

### 2.4. Molecular Dynamics Simulations

NAMD engine (v2.15) was used to simulate all systems [41] using the CHARMM36m force field [42, 33, 43]. Waters were explicitly treated by the TIP3P model [44]. Periodic boundary conditions were set for all systems and an integration time-step of 2 fs was used for all simulations. Long-range electrostatic interactions were computed with a grid spacing of 1 Å by applying the particle-mesh Ewald method [45]. The cutoff distance for non-bonded interactions was set as 12 Å. For soluble systems, energy minimization was performed for 10,000 steps followed by an NVT equilibration at 310 K for 250 ps. For transmembrane systems, a multi-step equilibration procedure was applied where force constants on protein and lipid atoms were gradually reduced over six cycles. Force constants applied to (i) the protein backbone atoms were gradually reduced from 10.0 to 0.1, (ii) the side chain atoms were reduced from 5.0 to 0.0, (iii) the membrane atoms were reduced from 2.5 to 0.0 over six cycles. First four cycles lasted 250 ps and the last two cycles lasted 500 ps. Production simulations were conducted for over 500 ns under constant temperature (310 K) and pressure (1 atm) in NPT ensembles using the Langevin thermostat and piston pressure method [46, 47, 48]. System details were given in supplementary Table 1.

### 2.5. Trajectory Analysis

Production trajectories were analyzed by pairwise root mean square displacement (RMSD) and fluctuation (RMSF) of Cα atoms using MDanalysis [49, 50] and in-house scripts. Solvent-accessible surface areas (SASA) and distance analysis were conducted using in-house scripts. Essential dynamics were extracted by principal component analysis (PCA) using Bio3D [51]. For visualization of structures and trajectories, VMD and ChimeraX were used [19, 52, 53, 54].

### 2.6. Prediction of Binding Free Energy

Binding free energy (ΔG) of all complexes were calculated as the average of 5 conformations taken from the last 20 ns part of the production simulations. The PRODIGY (PROtein binDIng enerGY prediction), which uses an empirical scoring function, was used to determine the binding free energy of the protein-protein complexes [31, 55, 56, 57].

## 3. Results

### 3.1. Molecular Modeling of All IL37 Complexes Involved in Anti-inflammatory Response

We initially explored rigid and flexible docking approaches to generate the binary complex structure of IL37-IL18Rα. However, neither of the methods provided reliable models that align with the experiments. Specifically, rigid-body docking produced complex models wherein either IL37 interacted with the IL18Rα through a surface excluding the third Ig-like domain (D3) or had a misaligned β-trefoil structure with the IL18 in the receptor complex (Fig. S1a). We also carried out flexible docking restricting the search space to amino acids at the interface of IL18-IL18Rα and keeping the N-terminal loop of IL37 (residues 49-57) fully flexible. Although flexible docking resulted in more relevant binding poses than the rigid-body models wherein the N-termini of both cytokines partially overlap, their trefoil cores still did not align well with each other (Fig. S1b). We also used AF2 multimer to generate the IL37-IL18Rα and IL37-IL18Rα-IL1R8 complexes. AF2 multimer predictions, as well, did not result in models compliant with the experimental findings as they showed that IL37 in a different orientation than IL18 in the binary complex and binds to the exterior region of the receptor leaving the IL18 binding site accessible in the ternary complex (Fig. S1c). The confidence scores of AF2 predictions including pLDDT, PAE and ipTM further implied a low accuracy for the interface region of both models (Fig. S1c). Hence, we noted that docking along with AF2 multimer predictions failed to produce experimentally relevant models of the IL37-IL18Rα and IL37-IL18Rα-IL1R8 complexes.

To generate an IL37-IL18Rα model consistent with experimental findings, we employed homology modeling based on the structural similarity between IL37 and IL18 (Fig. S2a). Superimposing the IL37 monomer onto IL18 in the binary PDB complex (3wo3) resulted in an optimal alignment of the β-trefoil fold and a short 3_10_ helix present in both structures, while a few short surface helices and loops differed between the two cytokines (Fig. S2b). The homology model exhibited a relatively low RMSD of 0.97 Å for the core (∼70 residues) and 4.27 Å for the entire chain (Fig. S2b). This model revealed an intermolecular salt bridge with the D3, an interaction also observed in the IL18 binary complex. Specifically, the interaction between K89 (UniProt: Q14116, K53 in PDB ID: 3wo3) of IL18 and E253 of IL18Rα is complemented by K53 in IL37 within the IL37-IL18Rα complex (Fig. S2b). Despite the low sequence identity between the two cytokines (∼20%), these two lysines in IL18 and IL37 maintained a similar spatial orientation towards E253 of the receptor (Fig. S2b), even though K53 of IL37 is located in the N-terminal region while K89 of IL18 is situated in the β1 strand. Taken together with all modeling efforts, the homology model of the IL37-IL18Rα complex provided credible structural representations that aligned well with experimental observations, outperforming docking and AF2 multimer models (Figs. S1-2).

Relying on IL18 homology, two distinct IL37-IL18Rα binary complexes were created using different IL18Rα structures from binary and ternary IL18 receptor complexes (PDB IDs: 3wo3, 3wo4) (Fig. 1). To assess the influence of the N-terminus of IL37, we further constructed binary complexes with distinct IL37 mature forms (49-206, 53-206, and 57-206). Additionally, the ternary complexes of IL37 involving IL1R8 and IL18Rβ as co-receptors were generated. Essentially, IL18Rβ containing complexes were used as negative controls as they were not reported experimentally. Our modeling extended to protein-protein complexes implicated in IL37’s receptor-independent anti-inflammatory actions, including Smad3 and IL18BP. As controls, we incorporated binary and ternary signaling complexes of IL18 from the PDB. Finally, full-length binary and ternary complexes of IL18 and IL37 were constructed. A comprehensive overview of the studied complexes was presented in Fig. 1 and Table 1. Together, these extensive structural models of IL37 and IL18 provide a foundation for elucidating the molecular role of IL37 in IL18 signaling.

**Figure 1:**
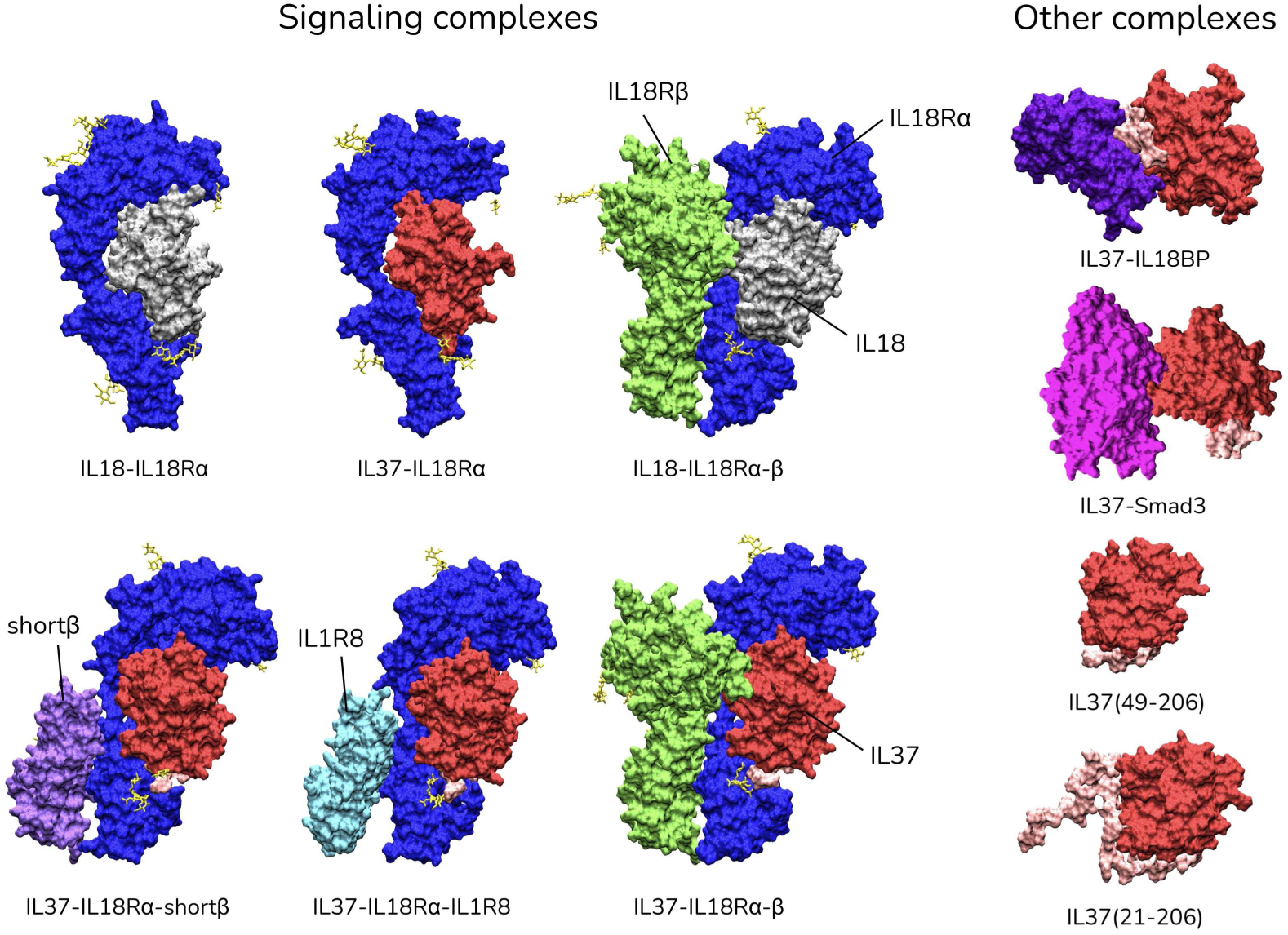
Overview of the PDB complexes of IL18 and IL37 complexes modeled in this study. Each subunit is represented in surface renders; IL37: red, IL18: grey, IL18Rα: blue, IL18Rβ: green/light purple, IL1R8: cyan, IL18BP: purple, Smad3: magenta. For monomeric IL37s, the N-terminal loop is colored in pink.

### 3.2. Soluble IL37 Complexes Display Higher Backbone Mobility Than IL18 Complexes

All IL37 complexes listed in Table 1 were analyzed in MD simulations. Homology models of the IL37 and IL18Rα produced some clashes at the N-terminus of IL37, as raised previously [15]. Since we checked the clashes prior to production simulations, minimization and equilibration steps resolved these clashes, reflecting that IL37 maintains the IL1 family binding mode in the IL18 receptor complexes without any steric clashes.

Pairwise Cα RMSD calculations were conducted to assess overall backbone mobility for the systems containing IL37 complexes (Fig. S3a) and IL18 complexes (Fig. S3b). For each system, pairwise RMSD calculations were made referencing the entire polypeptide chains (Fig. S3-ref:complex) and referencing the cytokine backbone (Fig. S3-ref:IL37/IL18). IL18-IL18Rα binary complexes showed almost no mobility in neither of the chains, albeit one of the IL37-IL18Rα models showed a high mobility for the IL18Rα with RMSD values exceeding 30 Å (Fig. S3a). For this system, three replicate simulations were conducted using different IL18Rα structures from the binary (3wo3) and ternary (3wo4) PDB complexes. Also, one of these simulations were elongated to ∼1 µs to further inspect the IL37 dynamics in the binary complex (Table 1). Albeit not as high as the first system, we noted a level of mobility for the receptor chains in these systems as well (Fig. S3a). We noted here that the receptor α complex formed by the shortest IL37 that completely lacks the N-terminal loop (57-206) showed an overall rigid backbone similar to the IL18 complex, underscoring the contribution of the N-terminal of IL37 to the receptor dynamics. In line with the RMSD analysis, the essential dynamics (ED) analysis suggested that IL37-IL18Rα complex covered a larger conformational space than the IL18-IL18Rα (Fig. S4). Notably, one of the highly mobile IL37-IL18Rα systems displayed a two-state conformation across PC1 scores (Fig. S4). The IL37 forms covering the regions 53-206 and 57-206 mostly showed PC1-PC2 score distributions similar to that of IL18-IL18Rα complex, implying relatively less mobile complexes in the shorter IL37 constructs. Accordingly, IL37-IL18Rα complexes with longer N-termini exhibited higher fluctuations in the D3 of the receptor, while the shorter forms (53-206 and 57-206) fluctuated less than this form for the entire complex (Fig. S5a), reflecting a more rigid complex structure for the shorter IL37 constructs.

Ternary complexes of IL37 also showed larger Cα RMSD values than that of IL18 (Fig. S3a-b). Particularly, the accessory receptors were highly dynamic in the IL37-IL18Rα-β and IL37-IL18Rα-IL1R8 complexes. However, we noted that the co-receptor in the IL37-IL18Rα-shortβ complex was mobile in the initial part of the simulation, it then achieved a relatively stable state. The ternary complex of IL18 also showed higher mobility compared to its binary complex. Nevertheless, the presence of a large disordered region of IL18Rβ (50-150), which is missing in the PDB structure (3wo4), could contribute to this increase in mobility (Fig. S3b). We also noted a lower level of fluctuations for the D3 in the ternary complexes (Fig. S5b). This effect is attributed to the presence of the accessory receptor that is docked to the C-terminal part of the receptor α, wherein the D3 lies.

We investigated the solvent accessible surface area (SASA) of IL18 and IL37 complexes (Fig. 2). Despite their similar size, IL37 had a larger surface area than IL18, especially for constructs with a longer N-terminal loop. Truncation of the N-terminus in IL37 (57-206) reduced its surface area to a level comparable to IL18 (Fig. 2a). Specifically, the surface area of IL18 ranged from 90-95 nm^2^, while that of IL37 (57-206) ranged from 86-95 nm^2^ and of IL37 (49-206) ranged from 93-105 nm^2^. Shorter IL37 constructs (53-206 and 57-206) had surface areas similar to or smaller than IL18. Fig. 2b highlights the N-termini of both cytokines in complex with IL18Rα, focusing on the 57*^th^* position of IL37. The IL37 N-terminus extends away from the cytokine surface along the β1-strand of IL18, creating an exposed loop. Mobility and exposure of this loop likely contribute to complex dynamics, explaining the differential responses of distinct IL37 mature forms. Overall, initial MD analysis showed that IL37-receptor complexes are more dynamic than IL18 complexes. The N-terminal loop of IL37 appears to be an important factor in determining complex dynamics, suggesting that shorter mature forms interacted more stably with the receptor α.

**Figure 2:**
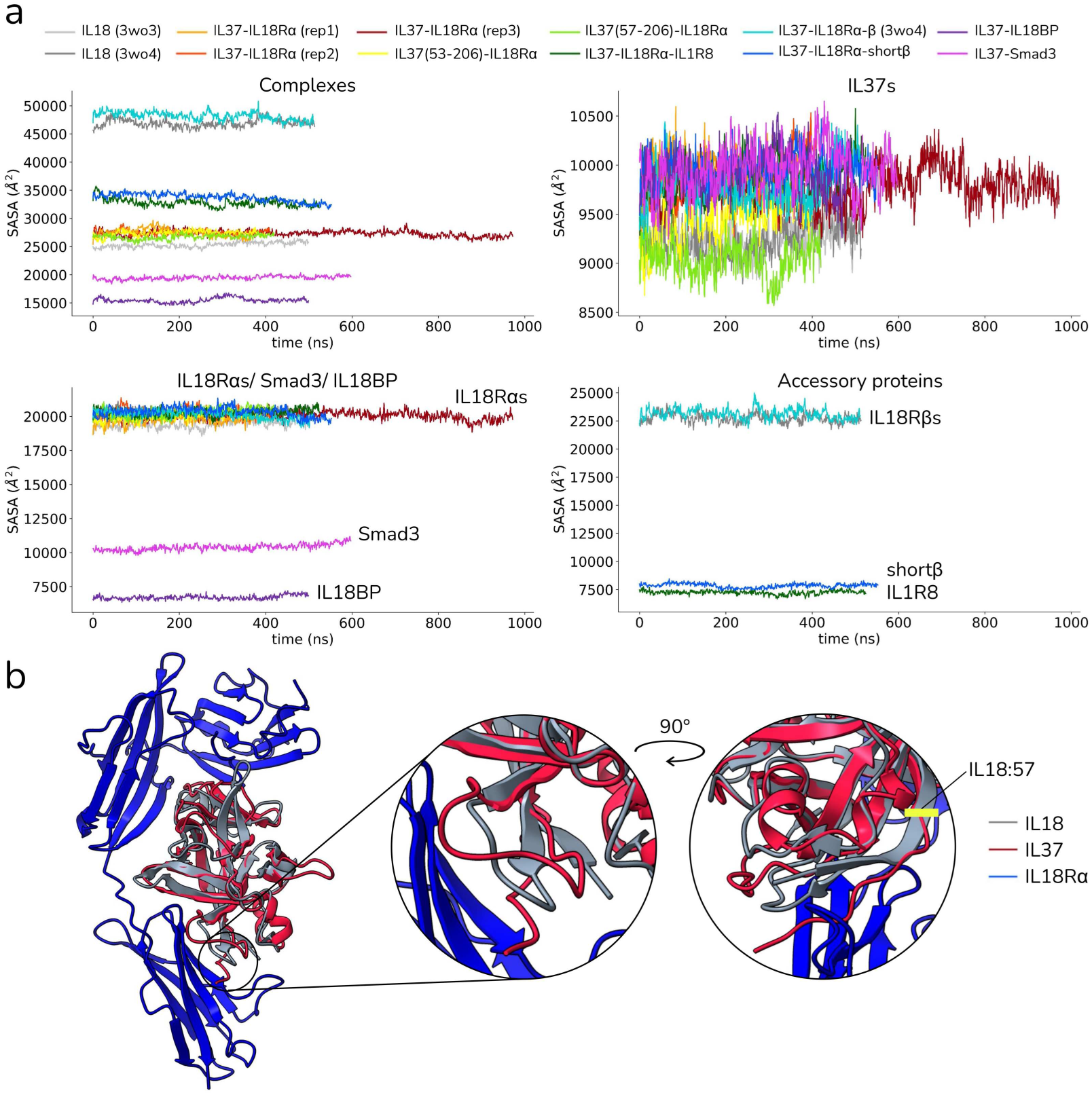
a) SASA of complexes and individual subunits. b) The homology model of IL37-IL18Rα complex is superimposed on the IL18-IL18Rα (PDB ID: 3wo3) complex. Close-up views illustrate the overlaid the N-terminal loop of IL37 on the β1-strand of IL18. The initial position of the N-terminal loop, specifically the 57*^th^* position, was highlighted in yellow.

For the other IL37 complexes formed by IL18BP and Smad3, despite neither of the complexes showed any dissociation, we noted that a dynamic interaction for both complexes (Fig. S3a). Specifically, the IL37-IL18BP complex, which was generated via homology modeling and shares the binding mode of IL18, exhibited a significant alteration in the initial homologous binding mode (supplementary Movie 1). A comparison of this modified binding pose with that of the IL18-IL18BP complex (PDB ID: 7al7) revealed a partial overlap between the binding interfaces of the IL18-IL18BP and IL37-IL18BP complexes.

To further assess the stability of the complexes used in our study, we performed binding free energy calculations for both IL18 and IL37 complexes by averaging 5 selected conformations from the last 20 ns of the simulations (Table 2). IL37 showed lower affinity towards the receptor α and IL18BP than did IL18 over binary complex trajectories and 7al7 structure. Although not statistically significant, binding free energy predictions indicated a marginal influence of the N-terminus of IL37 on receptor binding.

**Table 2:** Binding free energy predictions.

| IL18 Complexes | $\Delta G^\dagger$ | IL37 Complexes | $\Delta G$ |
| --- | --- | --- | --- |
| IL18-Rα<br>(PDB: 3wo3) | $-13.16 \pm 1.02$ | IL37-Rα | $-11.94 \pm 0.95$ |
| | | IL37-Rα (2) | $-11.52 \pm 0.96$ |
| | | IL37-Rα (3) | $-12.48 \pm 0.82$ |
| IL18-Rα-Rβ<br>(PDB: 3wo4) | $-12.82 \pm 0.52$ | IL37(53-206)-Rα | $-10.34 \pm 1.34$ |
| | | IL37(57-206)-Rα | $-11.12 \pm 1.31$ |
| IL18-IL18BP$^\ddagger$<br>(PDB: 7al7) | $-10.4$ | IL37-IL18BP | $-7.48 \pm 0.31$ |
| | | IL37-Smad3 | $-7.34 \pm 1.01$ |

$^\dagger$ Binding free energy values were predicted by Prodigy [56], unit is in kcal.mol $^{-1}$ and is given as mean ± sd. $^\ddagger$ PDB structure is used for calculation.

### 3.3. IL37 and IL18Rα Engage Through a Single Section of the Receptor

To understand how the binding interface changes in these complexes, we calculated the interface solvent accessible surface area (iSASA) between each subunit, which is computed by subtracting the complex SASA from the sum of the individual subunit SASAs. IL18Rα has three C2-type Ig-like domains, two at the N-terminus (D1-D2) and one at the C-terminus (D3) (Fig. 3a). To determine which part of IL18Rα regulates IL37 binding, we also examined the binding surfaces of these individual domain combinations. Fig. 3b-d shows the changes in binding surface area for the entire IL18Rα (top panels) and for the D1-D2 and D3 (middle and bottom panels). The specific binding areas used in the iSASA calculations are highlighted in color on the right-hand side of Fig. 3b-d.

**Figure 3:**
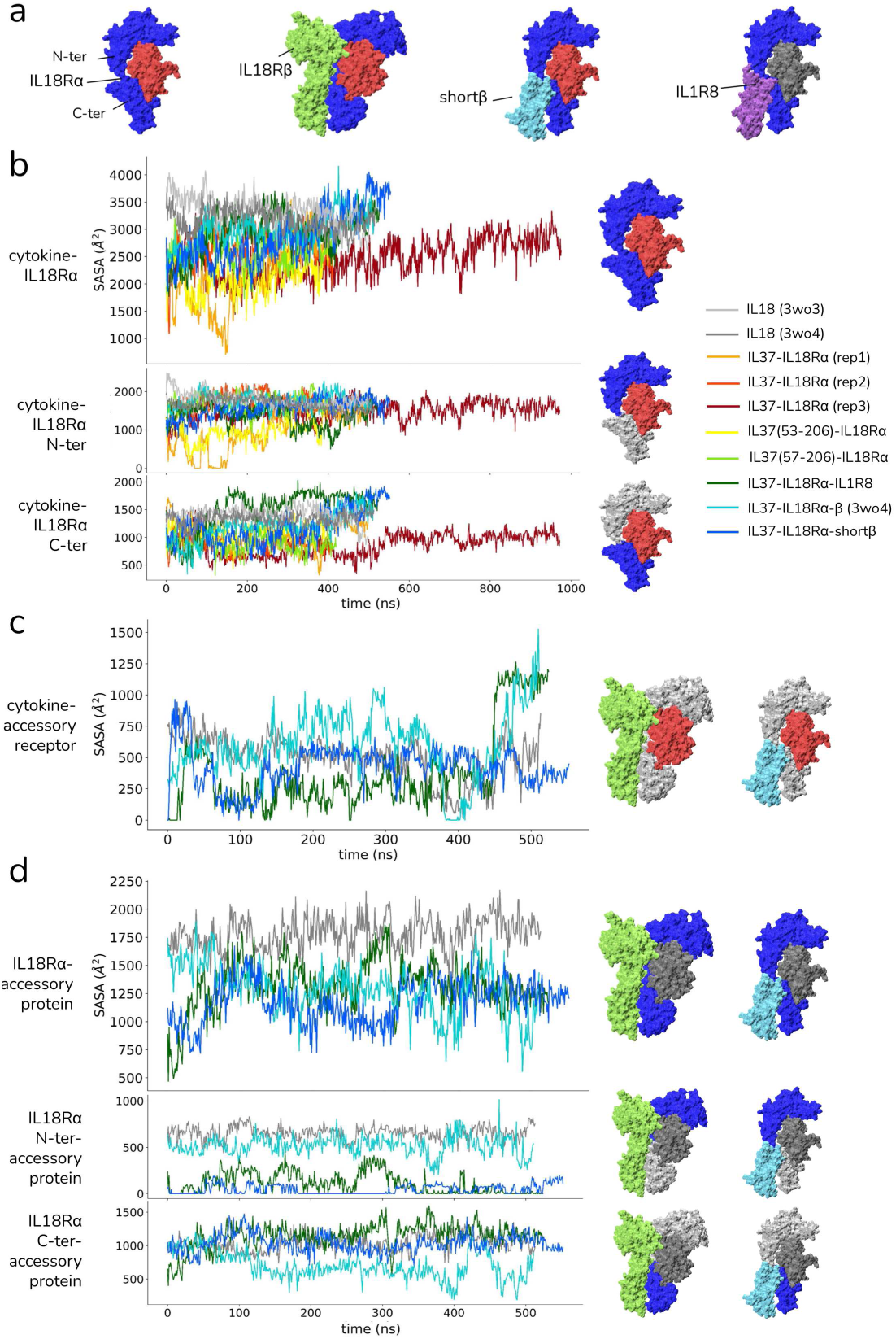
a) Receptor complexes were annotated by their subunits and the receptor α domains. b) Time-based variations in interface SASA (iSASA) between IL18Rα and the cytokines, IL18 or IL37. The top graph illustrates the iSASA for the entire IL18Rα, the middle graph depicts iSASA for the N-terminus (D1-D2) and the bottom graph for the C-terminus (D3) of IL18Rα. c) Changes in iSASA between the cytokines and the accessory receptors and (d) IL18Rα and accessory receptors. For b-d, the subunits used for iSASA calculations are shown in non-grey colors.

The buried surface area between IL18 and IL18Rα consistently fell within a range of 26-40 nm^2^ in both binary and ternary complexes (Fig. 3b). The interface area formed by either the N– or C-terminus of the receptor remained relatively unchanged, suggesting a stable interaction for IL18 with both segments of the receptor α. However, IL37 generally interacted with the IL18Rα through a smaller area. One particular IL37-IL18Rα complex exhibited a significant reduction in the buried area (Fig. 3b-orange). Upon examining the specific domains, this reduction was attributed to a loss of interaction between the D1-D2 corresponding to the N-terminus of IL18Rα (Fig. 3b-orange). The interface area reduced to zero at early stages of the simulation, indicating that this part of the complex had dissociated. The interaction between the N-terminus of the receptor and IL37 was re-established during the latter half of the simulation. In another IL37 complex, we observed a similar reduction in the interaction surface with C-terminal receptor, indicating loss of interaction with the C-terminus part of the receptor (Fig. 3b-dark red). This complex demonstrated the smallest interface area over a prolonged duration in the simulation (∼0.5 µs), although it achieved a more compact interaction in the latter half of the simulation. On the other hand, the interaction interface with the receptor’s N-terminal section remained stable for the prolonged simulation. The complex of IL37 spanning amino acids 53-206, also displayed a disrupted interaction with the N-terminal part of the receptor α, albeit to a lesser extent (Fig. 3b-yellow). Importantly, all disrupted interactions within the IL37-IL18Rα receptor complexes were ultimately restored in the extended portions of their simulations.

Ternary complexes involving IL37 exhibited the largest interface area with IL18Rα, reaching peak values towards the end of the simulations (Fig. 3b-cyan, blue, green). IL37 consistently achieved a larger interaction surface particularly with the D3 in all three complexes. Specifically, the ternary complex of IL37 with IL1R8 displayed distinct binding surface dynamics compared to the others (Fig. 3b-green). In the IL1R8 ternary complex, IL37 rapidly increased its binding surface with the D3 early on and maintained this interaction, while the binding surface with D1-D2 was slightly reduced.

Fig. 3c illustrates the binding surface areas of IL37/IL18 and co-receptors in the ternary complexes. Compared to the interaction area with receptor α, the co-receptors engaged with a smaller surface area for both cytokines. Additionally, the interaction surface area between cytokines and co-receptors was highly dynamic across all ternary complexes. Specifically, the interface area varied from 0 to 15 nm^2^ for all complexes, including the IL18 ternary receptor complex. The IL37 ternary complex with short-IL18Rβ demonstrated behavior akin to the IL18 ternary complex, leading to a similar buried surface area. On the other hand, the IL37 ternary complexes that included IL1R8 and the full β receptors exhibited a notable alteration in the binding surface with IL37, suggesting possible shifts in co-receptor orientations in these IL37 ternary complexes, except for the one involving short-IL18Rβ.

In the final part of iSASA calculations (Fig. 3d), we also analyzed the buried surface area between receptors. The IL18Rα and IL18Rβ interacted through a relatively large surface area (15-21 nm^2^) in the ternary complex of IL18 (Fig. 3d-gray). The corresponding surface area was consistently smaller in all IL37 ternary complexes. Specifically, the full form of the IL18Rβ primarily due to disrupted interactions with the C-terminus of the receptors (Fig. 3d-cyan). IL18Rα interacted with short-IL18Rβ (Fig. 3d-blue) and IL1R8 (Fig. 3d-green) through a similar surface area early in the simulation, but subsequently formed a more compact interaction with IL1R8 around 300 ns. Short-IL18Rβ (blue) interacted with the N-terminal domain of IL18Rα through a highly restricted region, with iSASA values ascending to 0 Å. IL1R8 (Fig. 3d-green), in contrast, exhibited relatively low but dynamic iSASA values with the IL18Rα N-terminal domain. It is noteworthy that short IL18Rβ and IL1R8 dimerized with the D3 through a surface area similar to that of IL18. Conversely, a restricted dimerization surface between IL18Rα and IL18Rβ was observed in the ternary complex of IL37 with full-length IL18Rβ. We also calculated the number of atom contacts between these subunits (<5 Å) (Fig. S6). The contact number analysis showed results that closely mirrored the iSASA calculations verifying that the interaction surfaces between the subunits were characterized by a matching number of atom contacts. Notably, the number of contacts between IL37 and its co-receptors, IL1R8 and IL18Rβ, initially dropped to zero and then were re-established, suggesting a reorientation of these co-receptors.

Altogether, these results mainly showed that IL37 complex interactions were primarily maintained by one part of the receptor, either D1-D2 (N-terminus) or D3 (C-terminus). The IL37 binary complexes demonstrated a unilateral binding mode to the IL18Rα, while IL18 exhibited bilateral binding, consistently interacting with both the N– and C-terminus lobes of the receptor α. These observations indicated a more dynamic receptor complex formation by IL37 than by IL18, highlighting that. On the other hand, IL37 ternary complexes displayed a large variability in the complex organization as reflected by fluctuations in iSASA analysis (Fig. 3c-d).

### 3.4. N-terminal Loop of IL37 Mediates IL18Rα Interactions

We focused on the simulations of the IL37-IL18Rα and IL37-IL18Rα-IL1R8 that have experimentally characterized. For comparison, we also examined the IL18 signaling complexes. The dynamics of these complexes were analyzed in detail using 1D RMSD calculations and structural visualization of trajectories. 1D RMSD plots and representative snapshots at specific time points are provided in Fig. 4 for binary complexes and in Fig. S9 for ternary complexes.

**Figure 4:**
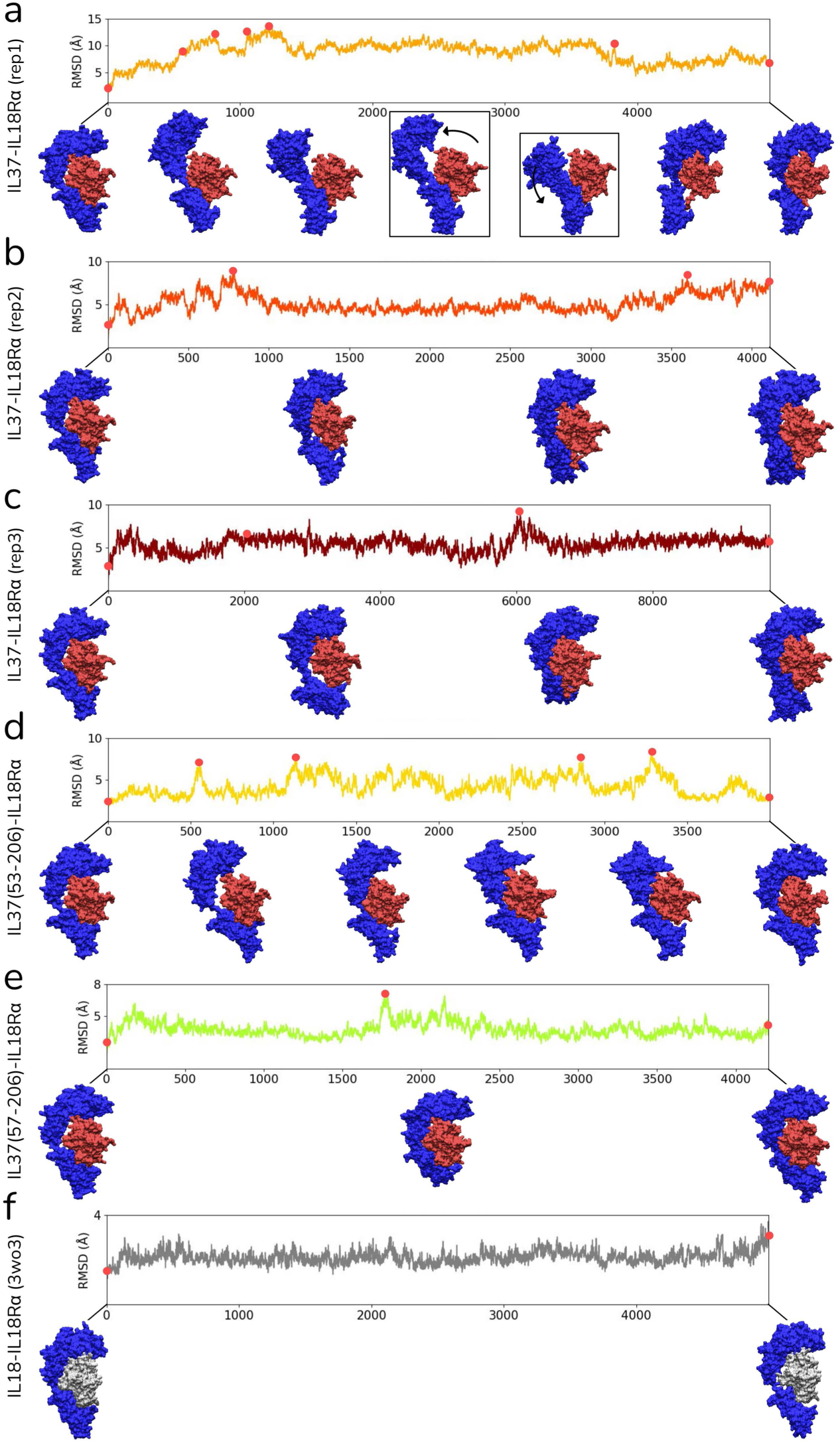
(a-f) 1D RMSDs were shown for the IL18 and IL37 complexes referencing the cytokine structure. Complex structures at specific time points with large RMSD were visualized.

We further focused on the highly dynamic simulation for the IL37-IL18Rα complex and observed that IL37 does not consistently maintain a bipartite mode of binding to IL18Rα as does IL18 (Fig. 4a, f). As illustrated in Fig. 4a, IL37 loses its interaction with either the N– or C-terminal lobe of the receptor α. IL18Rα terminal lobes hinged at the central loop connecting the D1-D2 to D3 oscillates around IL37, as the lost interactions were re-established for both receptor parts. We noted a contribution of the N-terminal loop of IL37 in the oscillation of receptor domains. In particular, one of the snapshots captured that the loop adopts a completely extended position on the C-terminus of the receptor towards the end of the simulation. Repeated simulations of this IL37-IL18Rα complex demonstrated similar variability in its interaction surface with the IL37-IL18Rα (Fig. 4b-e and Fig. S7). For all three simulations with longer loop forms, we reported that the loop initially positioned at the interface between IL37 and the D3 of IL18Rα adopts a compact conformation that wrapped around the IL37 core (Fig. S7). Subsequently, its conformation is transitioned to an extended, downward orientation, pushing the D3 away from IL37. This extended loop conformation induced a change in IL18Rα structure from a hook-like shape to an inverted L shape, reducing the contact area between IL37 and the D3 (Fig. 4a-c and Fig. S7). These observations implied that the IL37-IL18Rα complex with the same binding mode as IL18 is highly dynamic. In this binding mode, the cytokines interacted with the receptor α through two distinct surface regions, the N-terminal (D1-D2) or the C-terminal lobe (D3), whilst IL37 (49-206) tends to maintain only a single binding site with the receptor. Specifically, the N-terminal loop of this most studied isoform (49-206) affects the receptor interactions, disrupting intermolecular interactions by pushing the cytokine away from the D3 of the receptor towards its other domains.

The shorter loop of in the IL37 (53-206)-IL18Rα complex was also analyzed (Fig. 4d and Fig. S7). Yet, we did not observe a change as notable as in the IL37 (49-206) in the short N-terminal loop of IL37 in this system. Similarly, the IL37 (57-206) (Fig. 4e), when complexed with the receptor α, maintained the bipartite binding mode at all times as did IL18 (Fig. 4f).

In another simulation, we examined the dynamics of the IL37 monomer (49-206), concentrating on the conformations of its N-terminus (Fig. S3c). A trajectory of the loop (residues 49–57) of IL37 (49-206) mature form is shown in Fig. S8b. The monomer dynamics indicated that the N-terminal loop of IL37 explored highly accessible conformations that are separated from the IL37 core. Notably, the loop did not adopt a more compact orientation than its initial state through the simulation (Fig. S8b). Thus, the dynamic nature of IL37’s N-terminus in both monomer and complex forms is noted as an important factor mediating protein-protein interactions.

We further investigated the behavior of the longest mature form of IL37 (21-206) in MD simulations. One of the two replicate simulations revealed the N-terminal loop folding onto itself, creating a bulky extension on the IL37 surface (Fig. S8a-orange). The other simulation showed the extended loop wrapping around the IL37 surface, producing a globular form similar to shorter IL37 constructs (Fig. S8a-pink). Essentially, despite the added large loop at the N-terminus, the latter simulation, which displayed only a minor change in the receptor binding surface of IL37 (Fig. S8a-pink), suggests that an interaction with IL18Rα remains feasible in this state.

Our examinations revealed dramatically altered conformations of IL18Rα in complex with IL37, particularly in the D1-D2 (Fig. 4a). Two distinct IL18Rα conformations with the highest RMSD values were selected from the IL37-IL18Rα trajectories, and the IL37 dimer (PDB ID: 5hn1) was superimposed onto the monomeric IL37 on these complexes (Fig. 5). These complex snapshots reflected that such alternative IL18Rα conformations with altered orientations of the N– and C-terminus of the receptor could accommodate IL37 dimers without any steric clashes.

**Figure 5:**
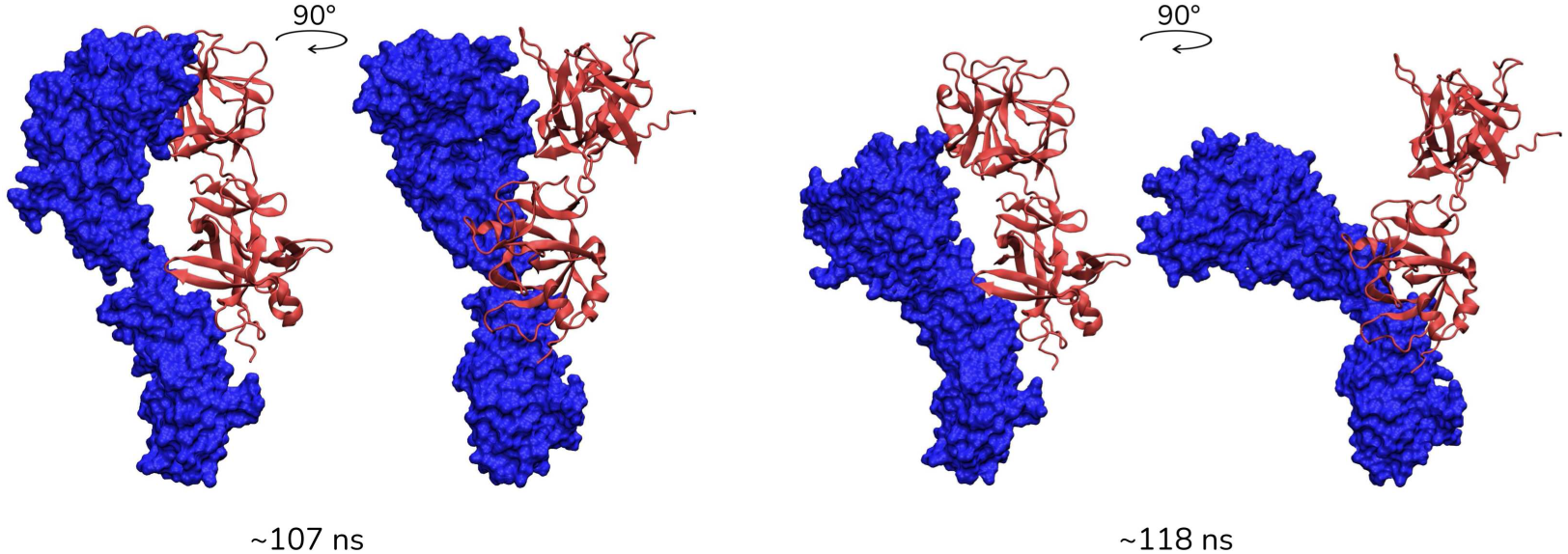
Two conformations extracted from one of the IL37-IL18Rα simulations were overlaid with the homodimer structure of IL37 (PDB ID: 5hn1). The extracted structures were framed in Fig. 4a.

### 3.5. A Glycan Linked to the Third Ig-like Domain of IL18Rα Shields the Cytokine Binding Site

IL18Rα has seven N-linked glycosylations in both binary and ternary IL18 signaling complexes. One of these glycosylations (GlyTouCan ID: G32152BH) is attached to N297 in the D3, located near the binding surface of the IL37 N-terminal loop (Fig. 6a). To investigate the glycan’s impact on the IL37 binding, particularly its N-terminal loop, we visualized the shortened trajectory of the loop and the glycosylated N297 (Fig. 6b) and measured the Cβ distance between residue K53 in the IL37’s loop and N297 in IL18Rα (Fig. 6c). A coupled movement of the N-terminal loop and the N297 was observed in the receptor α complexes containing the IL37 (49-206), showing a close location of the terminal loop position to the glycan. Distance distributions showed smaller variability in the the IL37 (49-206) while a higher variability in the spearation between glycan and loop in the shorter IL37 (53-206). Taken together with the reduced trajectories, we reported that the loop terminal residue and the glycan showed a coupled movement in the IL37 spanning 49-206 region, while this coupling was not present in the short form.

**Figure 6:**
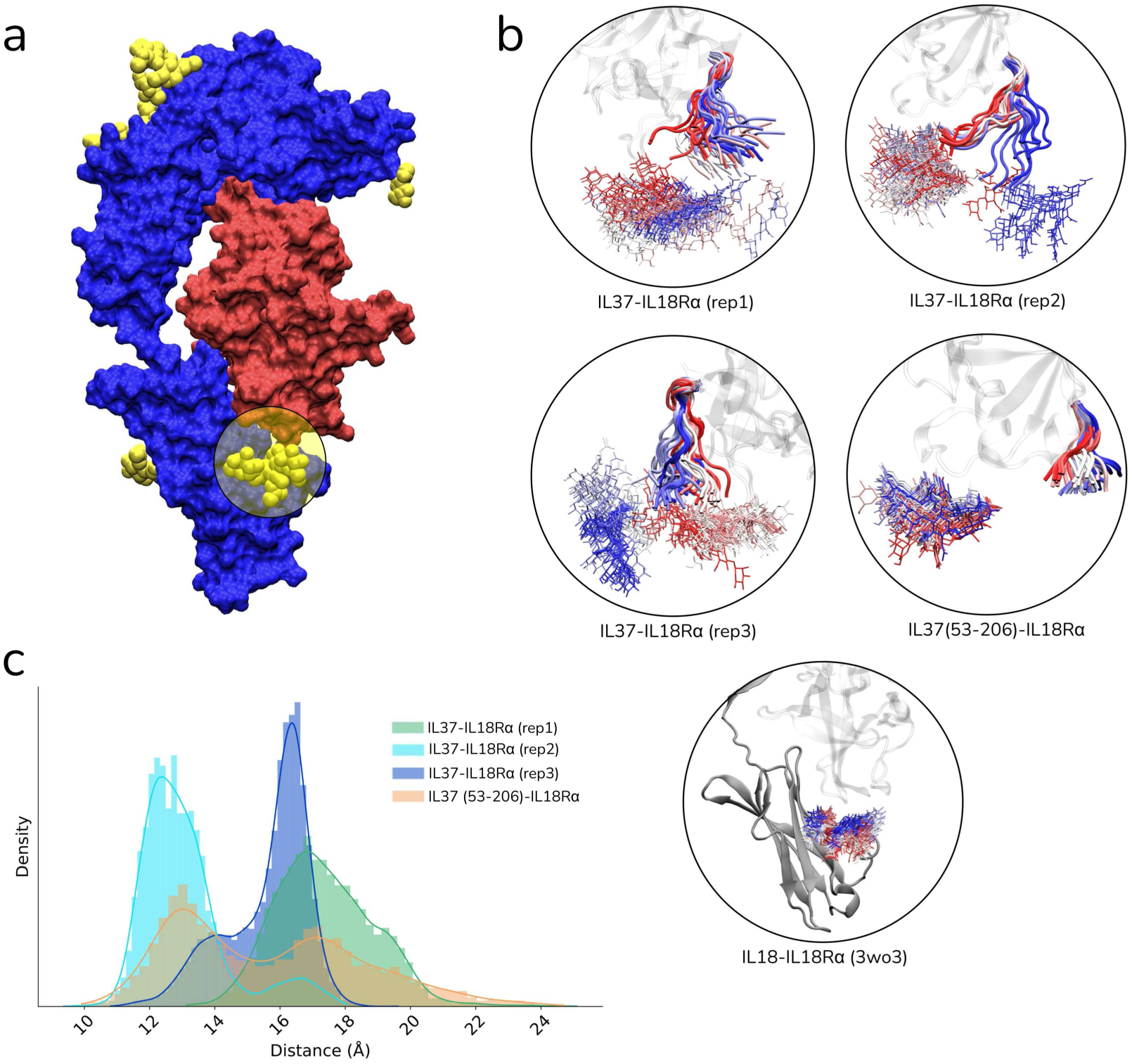
a) IL18-IL18Rα complex (PDB ID: 3wo3) is illustrated with glycosylations colored in yellow. b) Shortened trajectory of the glycoslyated N297 of the IL18Rα and the N-terminal loop of IL37 were shown. Red-white-blue color scale indicates simulation time-step. c) Cβ-Cβ distance distributions of the N297 of IL18Rα and P57 of IL37.

The glycated N297 acts as a shield of the cytokine binding site at the D3 of IL18Rα. This likely contributes to the conformational changes observed in the binary complexes (Fig. 4). The N-terminus of IL37 when faces the glycosylated N297 at the interface, we observed a distortion in the conformation of the D3, resulting in an inverted L-form of IL18Rα instead of a hook-like structure. This interaction sustains an open conformation of the N-terminal loop and IL18Rα. However, in shorter IL37 forms such as 53-206 and 57-206 or in IL18, the glycan and cytokine behaved more independent, reducing the glycan’s influence on the cytokine-receptor dynamics. In fact, the N297-linked glycan was found to be localized to a rather restricted area in the IL18-IL18Rα binary complex lacking the N-terminal loop and also in the IL37 (57-206)-IL18Rα complex, which mimics the behavior of IL18.

In ternary complexes, we did not observe any coupled movement between the cytokine’s N-terminal loop and the glycosylated residue N297 Fig. S9e. When a co-receptor dimerized with at least one tier of the receptor α, the glycated residue exhibited more confined dynamics, uncoupled from the loop dynamics of the cytokine, even in longer loops (Fig. S9a-c). These findings suggest that receptor dimerization stabilizes the receptor side of the interface but has little effect on loop dynamics. The cytokine loop remains highly mobile, independent of the glycan, similar to its behavior in single-receptor complexes.

### 3.6. Insights from the Full-length Transmembrane Complexes of IL18 and IL37

To construct the full-length signaling complexes for IL18 and IL37, we utilized AF2 to generate both binary and ternary configurations (Table 1). These complexes were then investigated by MD simulations. Fig. 7 shows pairwise Cα RMSD heatmaps, along with representative snapshots from the start and end of the simulations. The RMSD calculations were referenced using either the whole complex (largest heatmap) or specifically the extra-cellular (EC) or intracellular (IC) segments of the complexes (Fig. 7).

**Figure 7:**
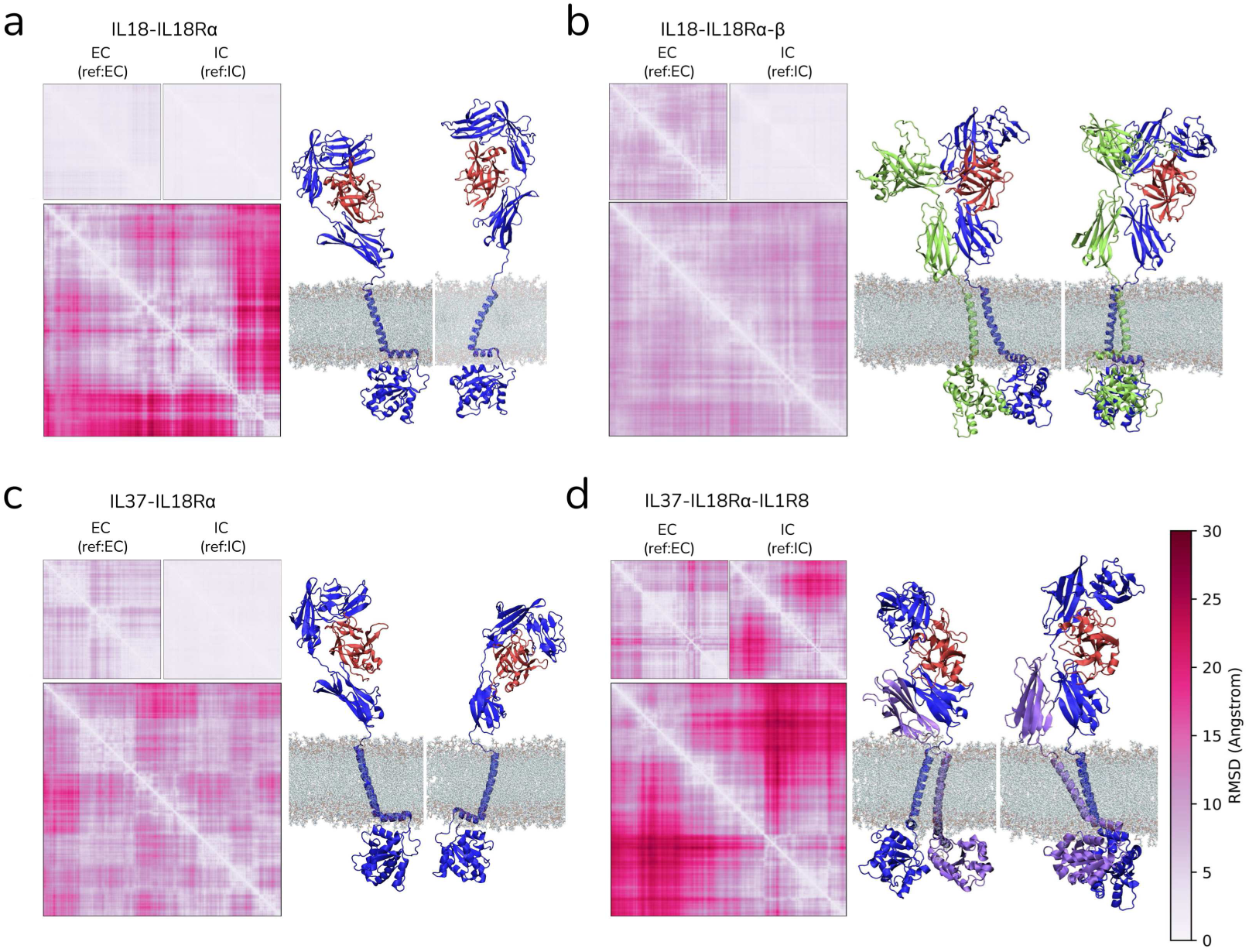
a-d) Pairwise RMSD heatmaps of the full-length complexes were shown. Largest RMSD heatmap was plotted referencing the entire complex, while top-left and top-right heatmaps were obtained referencing the EC (extracellular) and IC (intracellular) segments, respectively. Representative snapshots were extracted to illustrate the initial (left) and final (right) structures of the complexes. Color scale applied all panels.

The overall mobility of the full-length IL18-IL18Rα complex is attributed to the displacement of the transmembrane region, rather than the EC or IC segments, which maintained a stable RMSD profile (< 5 Å) (Fig. 7a). Initial and final structures from this trajectory demonstrate the flexibility of the transmembrane region leading to a 180*^o^* rotation of the EC part, while the IC region’s TIR domain remained unchanged. Similarly, the full-length ternary IL18 signaling complex displayed a stable RMSD profile for the IC region, with only slight mobility in the EC section (Fig. 7b). The TIR domain dimerization in the IC region kept its initial AF2-multimer predicted conformation throughout the simulations.

The full-length binary IL37 complexes were more mobile than IL18, aligning with the soluble complex simulations, showing higher mobility in the EC segment compared to the IC segment. This is shown by the first and last snapshots of the complex structure, where IL37 lost most contacts with the D3 (C-terminus) of the receptor (Fig. 7c), unlike IL18 complexes (Fig. 7a). For the ternary IL37 complex, we utilized the IL37-IL18Rα-IL18R1 complex based on the previous experimental data [11]. This complex exhibited a unique organization of the TIR domains in the IC section. Additionally, the Ig-like domain orientation of the IL1R8 was more intact than in the soluble complex (Fig. S9), indicating the transmembrane region’s role in maintaining receptor dimerization outside the cell.

Based on the TmAlphaFold database [25], the elongated vertical helices shown in Fig. 7 were identified as transmembrane segments in all three receptors. Our findings revealed that the short amphipathic helix, positioned orthogonally to the long TM-helix, was embedded within the membrane in the binary complexes. The insertion of this short helix is likely to be due to its amphipathic nature and the bulky extracellular (EC) portion of the complex. In contrast, in the ternary complexes, the helix remained at the membrane periphery, emphasizing its role in the dimerization of the extracellular (EC), intracellular (IC), and TM regions of the receptors.

## 4. Discussion

IL1 family of cytokines, including both IL18 and IL37, adopt a β-trefoil fold that is characterized by twelve β-strands, four of which packed together forming one of the three pseudo-repeats of the trefoil [15, 58]. Leveraging their structural resemblance, we constructed models of the IL37-receptor complexes based on IL18 homology. These models ensured that the D3 of the IL18Rα is positioned at the interface, a previously known interaction [4]. Otherwise, docking and AF2 multimer predictions failed to capture the same binding mode (Fig. S1).

Comparison with related cytokines [11] supports the shared binding mode of IL37 and IL18 with IL18Rα. Both IL37 and IL1RA act as receptor antagonists for IL18 and IL1β, inhibiting the proinflammatory signaling cascade [4, 59]. The PDB structures 1itb and 1ira represent IL1β-IL1R1 and IL1RA-IL1R1 complexes, with IL1RA being an IL1 receptor antagonist [59]. Superimposing IL1RA onto IL1β shows a backbone RMSD of 0.85 Å for the core and 3.48 Å for the entire chains. Our IL37-IL18Rα complex shows similar RMSD values of 0.97 Å and 4.27 Å for the core and full-length structures (Fig. S2b). Thus, the IL1β and IL1RA structural relationship further validates the predicted IL37 binding mode in our models.

In an earlier sequence alignment, K124 of IL37 was thought to correspond to K89 in IL18, potentially implicating them in IL18Rα binding [11]. Structural alignment of our modeled IL37 complexes with the IL18 complexes, however, showed another lysine (K53 in IL37) overlapping with K89 in IL18, both interacting with the E253 of IL18Rα (Fig. S2b). Given this possibly conserved salt bridge interaction with the D3 of IL18Rα, the K89 in IL18 is likely to be functionally countered by K53 in IL37, but not by K124 as proposed previously [11]. Similarly, the same sequence alignment suggested that E35 of IL37 corresponded to E42 in IL18 [11]. Nonetheless, in both the PDB structures of IL37 (PDB IDs: 5hn1 and 6ncu) and in our models, IL37 lacks a portion of its N-terminus, including E35 [60, 15]. Although sequence homology may degrade over time, structural homology often persists across extensive evolutionary periods [61, 62]. Consequently, structural alignment is likely to more accurately reflect the conserved binding mechanisms in comparison to sequence alignment, especially given the low sequence similarity between IL18 and IL37 (∼20%).

The monomeric IL37 exhibits greater anti-inflammatory capabilities compared to the dimeric form [15, 60, 63]. The reduced anti-inflammatory function of the dimeric form has been structurally attributed to the significant clashes between the IL37 dimer and the D1 and D2 of the IL18Rα [15]. Simulations unraveled significantly altered conformations of IL18Rα in complex with IL37, particularly in the D1-D2 (Fig. 4a, supplementary Movie 2). Overlaying the IL37 dimer onto the monomeric IL37 in complex with IL18Rα indicated that the alternative IL18Rα conformations can accommodate IL37 dimers without any clashes or that monomeric IL37 might dimerize on the IL18Rα (Fig. 5). On another note, the large displacement of the N-terminus of the receptor α, that involves a tilt of ∼90^◦^ around the interconnecting loop of two termini lobes, implies that recruitment of the accessory receptor, IL18Rβ, would be infeasible. Specifically, the Ig-like domain dimerization of the receptors, α and β, at the C-terminus would most likely fail if these tilted conformations were adapted by IL18Rα. To these ends, these alternative IL18Rα conformations ground a basis for the observation that IL37-IL18Rα complex recruits the orphan receptor of IL1R8, which has a single Ig-like domain, rather than IL18Rβ consists of 3 domains. Given the change in the relative orientations of the N– and C-terminal lobes of the receptor α, the IL37 binary complex maintains a ternary complex with an intact receptor β, but it could with a shorter receptor that possesses only single domain.

Our findings confirm the involvement of the N-terminal loop of IL37 in modulating the dynamics of specific receptor α complexes, indicating that longer mature forms may reduce receptor complex stability. While this conclusion aligns with experimental studies underscoring the importance of the N-terminal of IL37 [60], our investigation further elucidates the interaction between the N-terminal loop of IL37 and the glycosylated receptor residue, N297. The glycosylation at N297 has been shown to contribute to the binding of IL18, as its mutation to Q markedly diminished the binding affinity of IL18 [5]. Our simulations indicated that N297, with high-mannose glycosyl chains, restricts the cytokine binding site at the D3, functioning as a steric blockade. This restriction operates as a molecular sieve, permitting stable receptor α interactions at the D3 site, exclusively for cytokines of certain sizes. Given that the IL1 family binding mode maintains their N-termini at this locale, this glycan shield preferentially facilitate the binding of specific isoforms of the cytokines while impeding those with longer or bulkier N-terminal regions.

Proximity ligation assays and FRET analysis have previously demonstrated the co-localization of IL37 with IL18Rα and IL1R8 within sub-40 nm proximities, indicative of a ternary complex formation comprising IL37, IL18Rα, and IL1R8 [64]. Thus, our comprehensive list of IL37 complexes included the IL37-IL18Rα-IL1R8 ternary complex. Nevertheless, this complex exhibited significant mobility, particularly in the initial pose of IL1R8 (Fig. S9). In contrast, the ternary complex that includes a complete trans-membrane region demonstrated a more stable complex backbone (Fig. 7 and Fig. S3a), illustrating the stabilizing impact of incorporating transmembrane and TIR domains into the modeling. Nevertheless, it should be noted that, apart from the ternary signaling complex of IL18, simulations of the IL37 ternary models yielded inconclusive results, highlighting the need for further experimental investigation.

Consistent with the binary complex, it was hypothesized that the IL37-IL18BP interaction would utilize the same binding interface as the IL18-IL18BP complex [4]. While this shared binding of IL18 and IL37 to IL18BP would contradict the anti-inflammatory function of IL37 on IL18 signaling—by diminishing the neutralizing capacity of IL18BP on IL18 binding—it was proposed that IL37-IL18BP might inhibit the formation of the ternary receptor complex by interacting with and depleting the accessory receptor of IL18R β [4]. Our findings indicated that the IL37-IL18BP complex, sharing the same binding mode as the IL18-IL18BP (PDB ID: 7al7), did not retain this binding mode (supplementary Movie 1), instead transforming into a different complex with a partially overlapping interface with the IL18-IL18BP complex. We hypothesize that an alternative mechanism, potentially regulated by the N-terminal loop of IL37 or another protein partner, may be involved.

In this study, we mainly focused on the regulation of the N-terminal loop of IL37 towards receptor binding because IL37 has an extended N-terminal loop compared to other cytokines of IL1 family, while they all have a β-trefoil core. Besides the N-terminal loop, the C-terminal loop of IL37 was also thought to contribute to IL37-receptor dynamics [60, 15]. Further research are required to shed light on the regulation of the C-terminal loop of IL37 in receptor dynamics, which serves as one of the future studies on the anti-inflammatory effect of IL37.

The IL37 complexes generated herein consistently showed higher mobility compared to the IL18 complexes. This increased dynamic behavior may explain IL37’s lower binding affinity with the primary partners of IL18 [12, 4]. Despite extensive research spanning over a decade into IL37’s anti-inflammatory properties, the absence of experimental structures of IL37 complexes might be attributable to its low affinity binding with receptors and/or other partners. The low affinity and pronounced dynamism of IL37 in its protein-protein complexes could complicate the experimental determination of complex structures. Detailed computational analyses, such as those presented herein, may provide insights into these experimentally challenging and versatile structures, thereby guiding future experimental efforts. Hence, we underscore the pivotal role of the interaction between the N-terminal loop of IL37 and the glycosylation at N297 of IL18Rα in fostering stable IL37 interactions *in vitro*.

## 5. Data Availability Statement

All IL37 complexes generated in this study are available at the https://doi.org/10.5281/zenodo.13822020.

