## Supplementary File 1 for "Computational Modeling of the Anti-Inflammatory Complexes of IL37"

### Supplementary Information
Computational Modeling of the Anti-Inflammatory Complexes of IL37

Inci Sardag[a], Zeynep Sevval Duvenci[b], Serkan Belkaya[c], Emel Timucin[b,d]

[a] *Bogazici University, Department of Molecular Biology and Genetics, Istanbul 34342, Turkey,*
[b] *Acibadem University Institute of Health Sciences Department of Biostatistics and Bioinformatics Istanbul 34752 Turkey*
[c] *Bilkent University, Department of Molecular Biology and Genetics, Ankara 06800, Turkey,*
[d] *Acibadem University Institute of Health Sciences Department of Biostatistics and Bioinformatics Istanbul 34752 Turkey*

---

#### List of Tables

1 Details of MD Systems . . . . . . . . . . . . . . . . . . . . . . . 4

#### List of Figures

1 (a) Clusters obtained from the rigid-body docking were visualized (b) Top cluster of the flexible docking analysis was visualized along side the PDB complex of IL18-IL18R$\alpha$. (c) AF2 predictions of IL37-IL18R$\alpha$, IL37-IL18R$\alpha$-IL1R8, and IL37-Smad3 complex structures were shown according to pLDDT confidence score coloring (yellow: plDDT<60, green: plDDT<70, cyan: plDDT<80, blue: plDDT<90). PAE heatmaps were also given at the bottom of the structures. Blue-orange-white color scaling represents error scores from 0 to 30. . . . . . . . 6

2 a) Binary complexes of IL18 (PDB ID: 3wo3) and IL37 formed by superimposition were shown. b) Monomeric forms of IL18 and IL37 were displayed with labeled helical differences. In close-up views, interaction between K53 of cytokines and E253 of IL18R$\alpha$ was shown. c) Binary complexes of IL1$\beta$ (PDB ID: 1itb) and IL1 receptor antagonist (IL1RA) (PDB ID: 1ira) were shown. d) Monomeric forms of IL1$\beta$ and IL1RA were displayed with labeled helical differences. . . . . . . . . . . . . 7
3 Pairwise RMSD plots were plotted for the C$\alpha$ atoms of (a) IL37 complexes, (b) IL18 complexes, (c) IL37 monomers. For complex structures, RMSD plots were created for the complex wherein the reference structure was complex, for the cytokine (IL37/IL18) wherein the reference structure was cytokine and for the partners wherein the reference structure was cytokine. For IL37 monomers, RMSD plots were plotted for the entire cytokine when aligned on itself, for the cytokine core when aligned to the core region (57-206) and for the N-terminal loop when aligned to the core. . . . . . . . . . . . . . . . . . . 8
4 Score plots for the first two PCs and scree plot for the explained variance were shown for all simulations. RWB color scale indicates the simulation time. . . . . . . . . . . . . . . . 9
5 RMSF of (a) cytokines and IL18R$\alpha$s, (b) cytokines, IL18R$\alpha$s and accessory proteins (c) cytokines and IL18BP or Smad3 were shown when each structure segment was aligned on itself. Accessory receptors in the ternary complexes were aligned to the single Ig-like domain at their C-terminus. . . . . . . . . . . 10
6 Left plots shows the number of atom contacts at the interfaces of binary, ternary and receptor-independent complexes and right plots show the number of unique amino acids involved in these contacts. . . . . . . . . . . . . . . . . . . . . . . . . 11
7 1D RMSD calculations of the N-terminal loop of IL37 in four binary complexes with two different mature forms of IL37 (49-206 and 53-206) were shown. Structures of the complexes at specific time points were also displayed at the bottom of the RMSD plots. The N-terminal loops were colored in pink. . . . 12

8 (a) (left) PDB complex of IL18-IL18R$\alpha$ (3wo3) is shown alongside two different mature forms of monomeric IL37, 49-206 and 21-206. The structures for the 21-206 form were obtained from two replicate simulations. N-terminal loops, 49-57 and 21-57, were colored to illustrate their conformation in the equilibrated structures. (b) Reduced trajectory of the N-terminal loop of the IL37 form covering 49-206. The initial conformation of the loop is colored in cyan while red-white-blue color scale indicates simulation time-step. . . . . . . . . . . . . . . . . 13
9 (a-d) 1D RMSD plots of cytokine (IL18/IL37) partners in four different ternary complexes were given. Representative snapshots from given time points were also shown at the bottom of the plots. (e) Reduced trajectory of the IL37's N-terminal loop and the glycosylated N297 were shown in three different ternary complexes. . . . . . . . . . . . . . . . . . . . . . . . . 14

#### 1. Tables

Table 1: Details of MD Systems

| | id | water | protein | $Na^+$-$Cl^-$ | Glycan | Membrane | xxyxz† | Duration |
|---|---|---|---|---|---|---|---|---|
| monomer | 1* | 2893 | 80442 | 76-78 | 0 | 0 | 96x96x96 | 500 ns / 550 ns |
| | 2 | 2498 | 27804 | 26-32 | 0 | 0 | 69x69x70 | 500 ns |
| soluble binary receptor complexes | 3 | 7271 | 155049 | 149-146 | 476 | 0 | 120x120x120 | 500 ns |
| | 4 | 7255 | 150426 | 142-149 | 482 | 0 | 120x119x119 | 450 ns |
| | 5* | 7289 | 158916 | 150-157 | 478 | 0 | 121x121x121 | 970 ns / 430 ns |
| | 6 | 7228 | 162378 | 154-159 | 478 | 0 | 122x122x122 | 400 ns |
| | 7 | 7157 | 162276 | 154-160 | 476 | 0 | 122x122x122 | 400 ns |
| soluble ternary receptor complexes | 8 | 12550 | 198072 | 192-186 | 656 | 0 | 131x131x131 | 510 ns |
| | 9 | 12511 | 198012 | 186-191 | 659 | 0 | 131x131x132 | 510 ns |
| | 10 | 9142 | 206505 | 195-203 | 572 | 0 | 132x132x132 | 550 ns |
| | 11 | 8967 | 206745 | 196-200 | 478 | 0 | 132x132x132 | 525 ns |
| full-length receptor complexes | 12 | 10779 | 203430 | 186-186 | 0 | 46230 | 119x119x237 | 380 ns |
| | 13 | 19381 | 346323 | 320-317 | 0 | 74504 | 148x149x246 | 450 ns |
| | 14 | 10742 | 203601 | 196-186 | 0 | 46230 | 120x120x237 | 350 ns |
| | 15 | 15708 | 162039 | 163-147 | 0 | 36180 | 108x110x239 | 400 ns |
| other complexes | 16 | 4143 | 76050 | 72-75 | 86 | 0 | 95x95x95 | 500 ns |
| | 17 | 5408 | 64974 | 61-69 | 0 | 0 | 91x91x91 | 500 ns |

System ids were taken from Table 1 in main text.
†Box dimensions were in Å units.
*Two replicate simulations were conducted for these systems.

Movie 1: Movie showing the MD trajectories of IL37-IL18BP complex formed in this study, aligned to IL18BP. On the side, static structures of 7al7 (IL18-IL18BP) and the last frame of the IL37-IL18BP complex are shown. Red, grey and blue color codes represent IL37, IL18 and IL18BP, respectively.

Movie 2: Movie showing the MD trajectories of one of the IL37-IL18R complexes together with the MD trajectories of IL18-L18R (3wo3) complex. Shows the altered conformation of the receptor in the IL37-IL18R complex. Red, grey and blue color codes represent IL37, IL18 and IL18R, respectively.

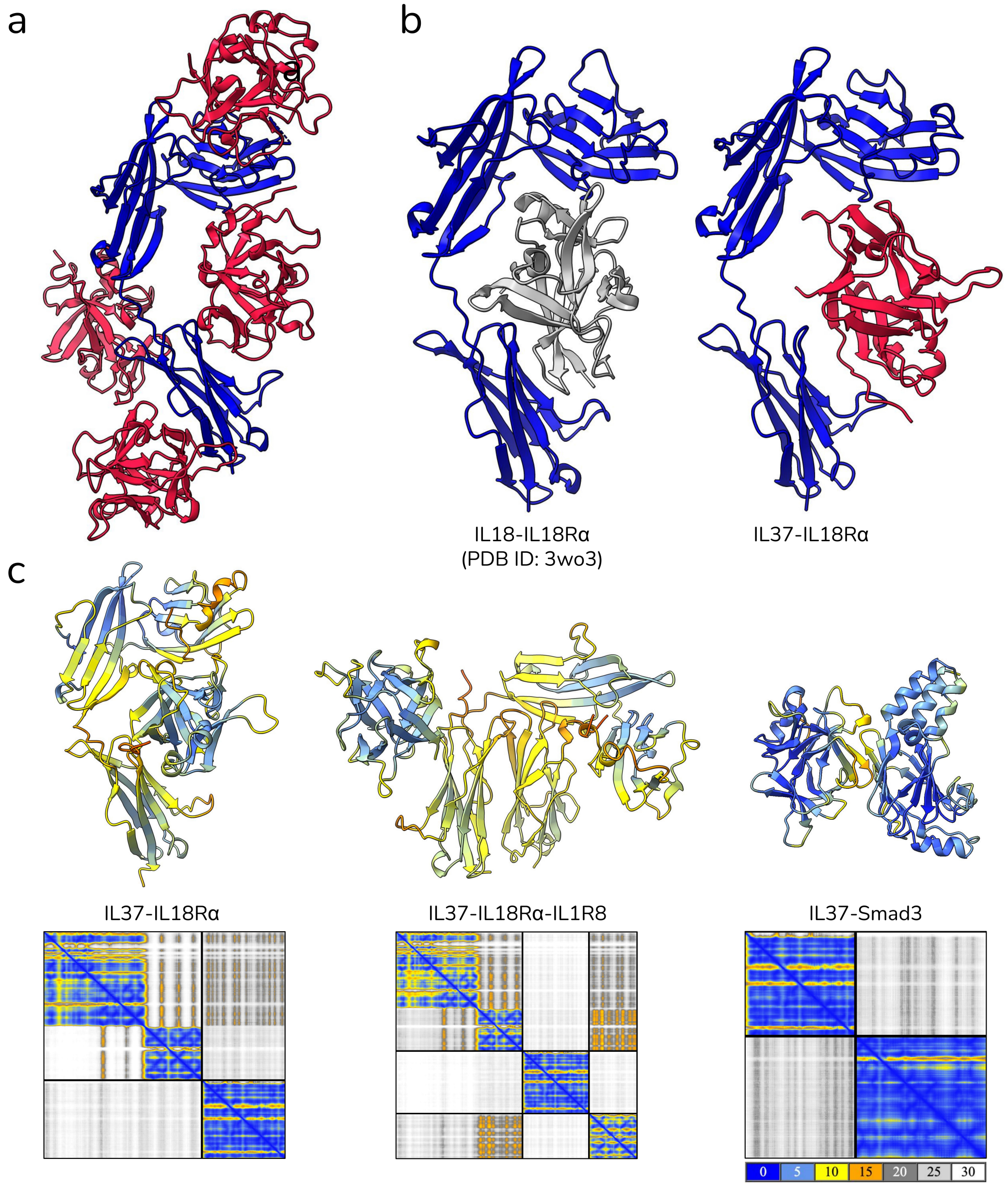


Figure 1: (a) Clusters obtained from the rigid-body docking were visualized (b) Top cluster of the flexible docking analysis was visualized along side the PDB complex of IL18-IL18R$\alpha$. (c) AF2 predictions of IL37-IL18R$\alpha$, IL37-IL18R$\alpha$-IL1R8, and IL37-Smad3 complex structures were shown according to pLDDT confidence score coloring (yellow: plDDT<60, green: plDDT<70, cyan: plDDT<80, blue: plDDT<90). PAE heatmaps were also given at the bottom of the structures. Blue-orange-white color scaling represents error scores from 0 to 30.

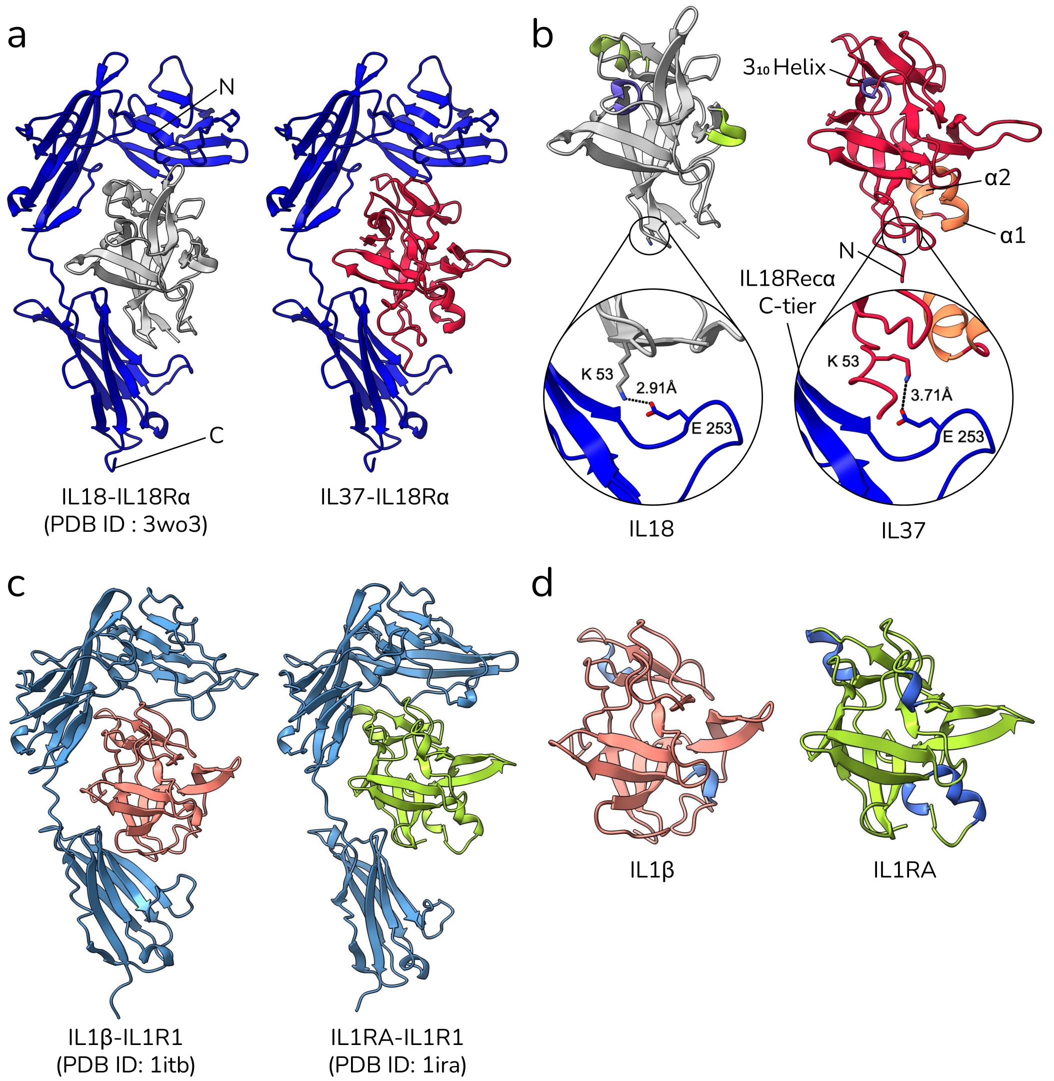


Figure 2: a) Binary complexes of IL18 (PDB ID: 3wo3) and IL37 formed by superimposition were shown. b) Monomeric forms of IL18 and IL37 were displayed with labeled helical differences. In close-up views, interaction between K53 of cytokines and E253 of IL18R$\alpha$ was shown. c) Binary complexes of IL1$\beta$ (PDB ID: 1itb) and IL1 receptor antagonist (IL1RA) (PDB ID: 1ira) were shown. d) Monomeric forms of IL1$\beta$ and IL1RA were displayed with labeled helical differences.

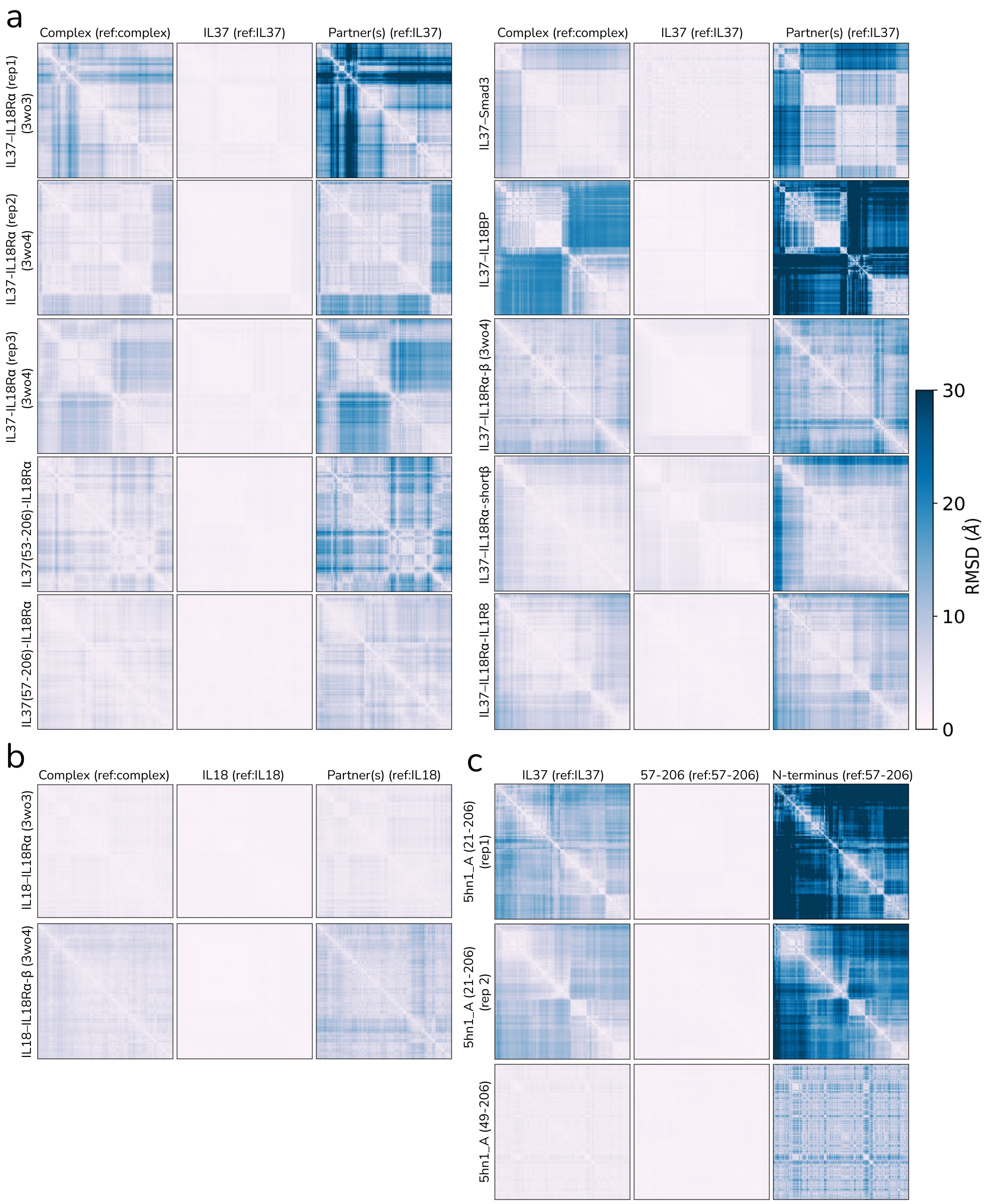


Figure 3: Pairwise RMSD plots were plotted for the C$\alpha$ atoms of (a) IL37 complexes, (b) IL18 complexes, (c) IL37 monomers. For complex structures, RMSD plots were created for the complex wherein the reference structure was complex, for the cytokine (IL37/IL18) wherein the reference structure was cytokine and for the partners wherein the reference structure was cytokine. For IL37 monomers, RMSD plots were plotted for the entire cytokine when aligned on itself, for the cytokine core when aligned to the core region (57-206) and for the N-terminal loop when aligned to the core.

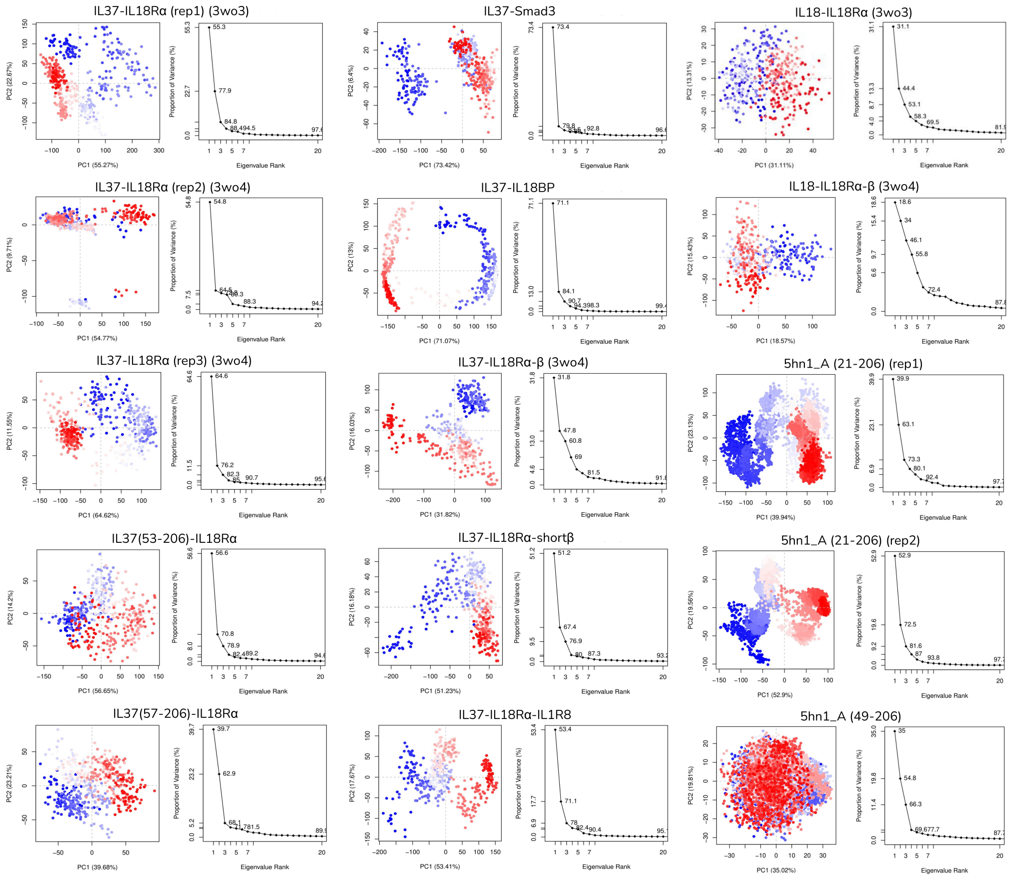


Figure 4: Score plots for the first two PCs and scree plot for the explained variance were shown for all simulations. RWB color scale indicates the simulation time.

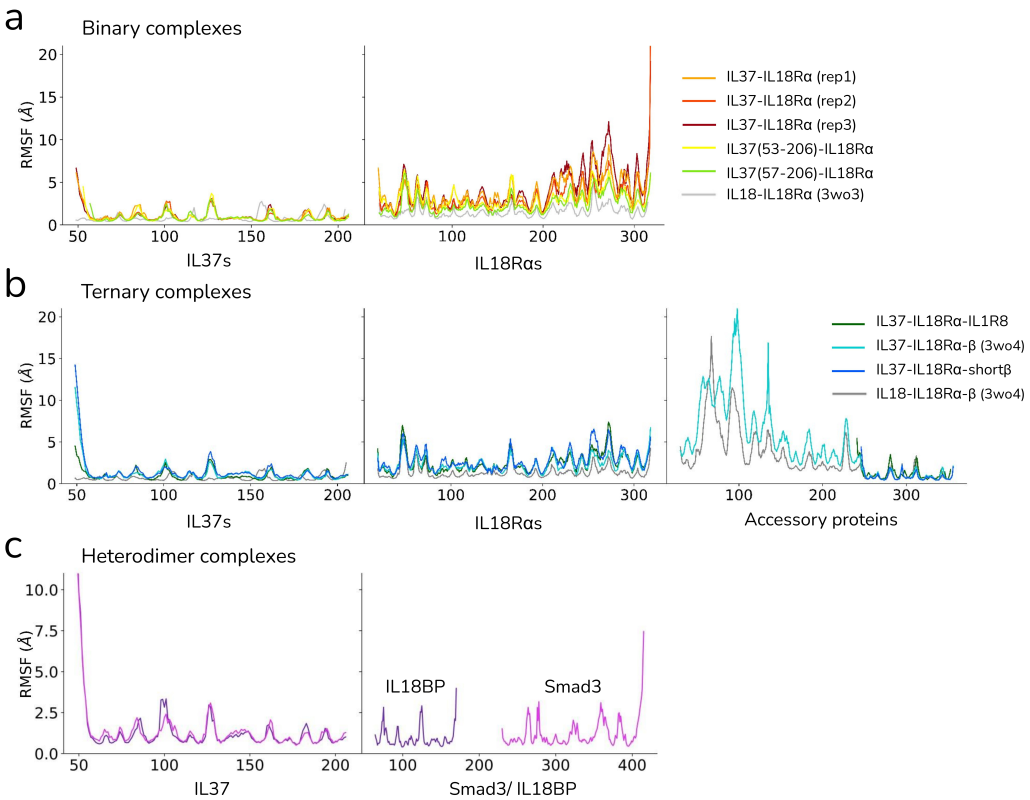


Figure 5: RMSF of (a) cytokines and IL18R$\alpha$s, (b) cytokines, IL18R$\alpha$s and accessory proteins (c) cytokines and IL18BP or Smad3 were shown when each structure segment was aligned on itself. Accessory receptors in the ternary complexes were aligned to the single Ig-like domain at their C-terminus.

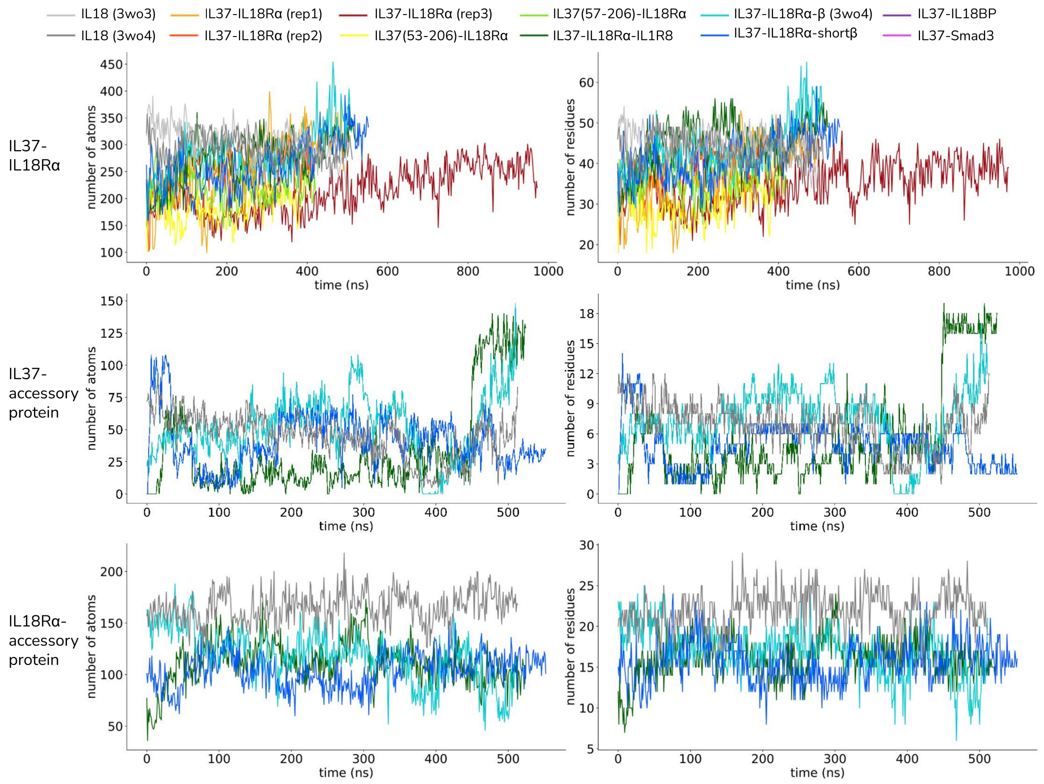


Figure 6: Left plots shows the number of atom contacts at the interfaces of binary, ternary and receptor-independent complexes and right plots show the number of unique amino acids involved in these contacts.

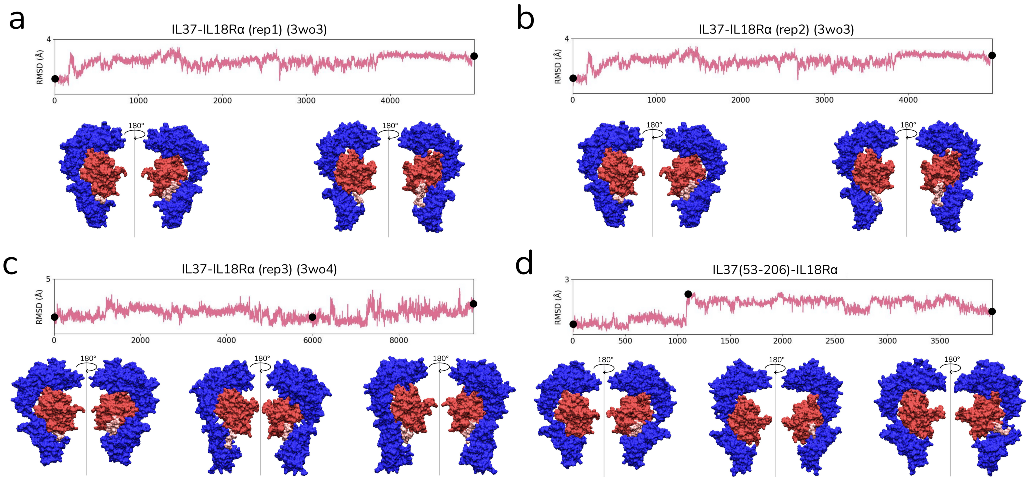


Figure 7: 1D RMSD calculations of the N-terminal loop of IL37 in four binary complexes with two different mature forms of IL37 (49-206 and 53-206) were shown. Structures of the complexes at specific time points were also displayed at the bottom of the RMSD plots. The N-terminal loops were colored in pink.

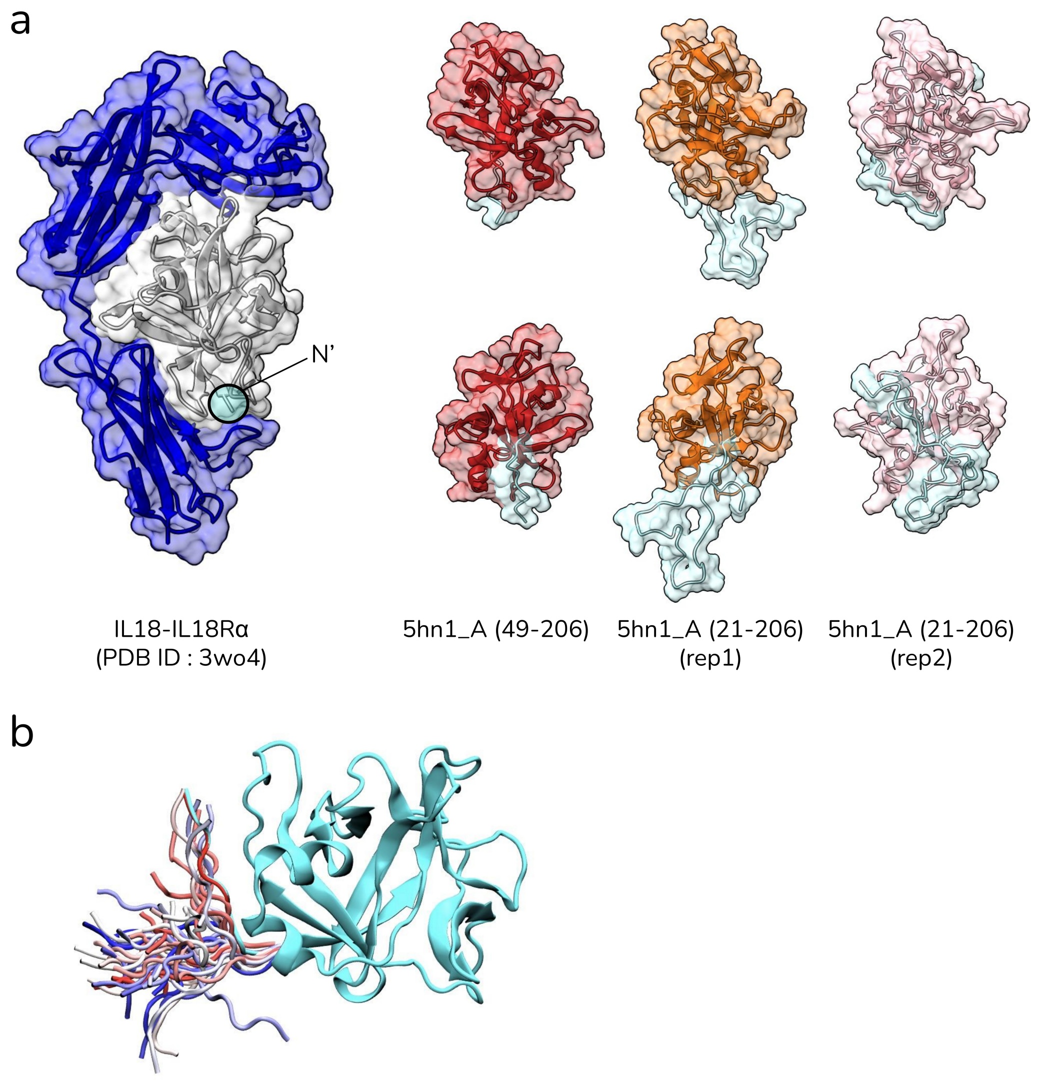


Figure 8: (a) (left) PDB complex of IL18-IL18R$\alpha$ (3wo3) is shown alongside two different mature forms of monomeric IL37, 49-206 and 21-206. The structures for the 21-206 form were obtained from two replicate simulations. N-terminal loops, 49-57 and 21-57, were colored to illustrate their conformation in the equilibrated structures. (b) Reduced trajectory of the N-terminal loop of the IL37 form covering 49-206. The initial conformation of the loop is colored in cyan while red-white-blue color scale indicates simulation time-step.

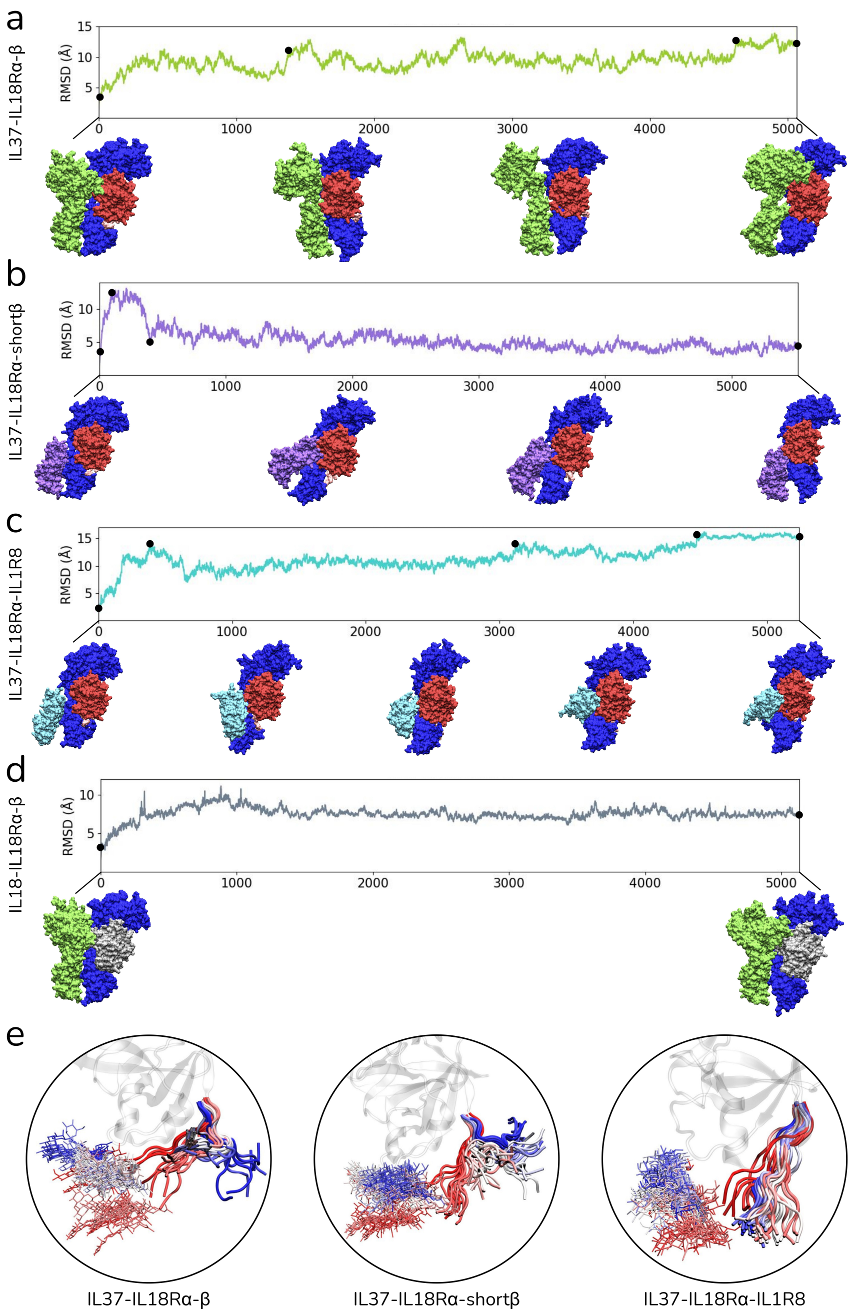


Figure 9: (a-d) 1D RMSD plots of cytokine (IL18/IL37) partners in four different ternary complexes were given. Representative snapshots from given time points were also shown at the bottom of the plots. (e) Reduced trajectory of the IL37's N-terminal loop and the glycosylated N297 were shown in three different ternary complexes.
